# Environmental and geographic data optimize *ex situ* collections and the preservation of adaptive evolutionary potential

**DOI:** 10.1101/2020.02.22.960989

**Authors:** Lionel N. Di Santo, Jill A. Hamilton

## Abstract

Maintenance of biodiversity, through seed banks and botanical gardens where the wealth of species’ genetic variation may be preserved *ex situ*, is a major goal of conservation. However, challenges can persist in optimizing *ex situ* collections where trade-offs exist between expense, effort, and conserving species evolutionary potential, particularly when genetic data is not available. Within this context, we evaluate the genetic consequences of guiding population preservation using geographic (isolation-by-distance, IBD) and environmental (isolation-by-environment, IBE) data for *ex situ* collections where provenance data is available. We use 19 genetic and genomic datasets from 15 plant species to (i) assess the proportion of population genetic differentiation explained by geographic and environmental factors, and (ii) simulate *ex situ* collections prioritizing source populations based on pairwise geographic or environmental distances. Specifically, we test the impact prioritizing sampling based on environmental and geographic distances may have on capturing neutral, functional or putatively adaptive genetic diversity and differentiation. We find that collectively IBD and IBE explain a substantial proportion of genetic differences among functional (median 45%) and adaptive (median 71%) loci, but not for neutral loci (median 21.5%). Simulated *ex situ* collections reveal that inclusion of IBD and IBE increases both allelic diversity and genetic differentiation captured among populations, particularly for loci that may be important for adaptation. Thus, prioritizing population collections using environmental and geographic distance data can impact genetic variation captured *ex situ*. This provides value for the vast majority of plant species for which we have no genetic data, informing conservation of genetic variation needed to maintain evolutionary potential within collections.

## Introduction

Genetic variation is fundamentally a prerequisite for adaptive evolution (Carlson et al. 2014). Consequently, to maintain species’ evolutionary potential, conservation often focuses on the preservation and maintenance of genetic variation. *Ex situ* collections provide one approach to preserve genetic diversity outside species’ native ranges. This includes extensive efforts to collect, preserve, and maintain variation across the range of different crop species, wild relatives, and rare or threatened species (Li et al. 2002; Westengen et al. 2013; Naredo et al. 2017). The Global Strategy for Plant Conservation (GSPC) aims to have at least 75% of endangered plant species preserved *ex situ* by 2020 and available for use in recovery or restoration (Target 8; https://plants2020.net/). While significant progress has been made, major gaps remain in the maintenance of genetic variation within collections (Sharrock et al. 2018). Consequently, *ex situ* programs designed to maintain genetic diversity are yet needed.

Traditionally, *ex situ* methods rely on either probabilistic equations (Brown & Marshall 1995; Lawrence et al. 1995), or stochastic resampling using pre-existing genetic datasets to optimize sampling efforts (Caujapé-Castells & Pedrola-Monfort 2004; Gapare et al. 2008). However, these approaches have limitations as they either require the availability of genetic data (population resampling strategy) or make ungeneralizable assumptions of within species population structure (probability-based strategy; Lockwood et al. 2007). More recently, simulation-based strategies have been developed and tested to guide sampling practices (Hoban & Schlarbaum 2014; Hoban 2019). Simulation-based approaches do not require previously published genetic datasets but enable realistic simulations of population structure using available estimates of population size and genetic connectivity. To overcome challenges associated with *a priori* data requirements, the use of surrogate data, such as environmental or spatial data, to estimate neutral and nonneutral genetic variation has received considerable attention (Guerrant Jr et al. 2013; Whitlock et al. 2016; Hanson et al. 2017). Empirical work has focused mainly on testing these data surrogates in preserving genetic diversity *in situ* or in wild populations (Whitlock et al. 2016; Hanson et al. 2017). However, using environmental and geographic data to optimize *ex situ* sampling could have substantial value to conservation.

Evolutionary processes have predictable impacts on the distribution of standing genetic variation, which may be used to guide *ex situ* collections. IBD or “isolation-by-distance” (Wright 1943) arises when gene flow between geographically distant populations is not enough to counteract the accumulation of genetic differences via genetic drift or following successive founder events during colonization (Slatkin 1993; Ledig 2000). In this way, IBD is a proxy for the relationship between pairwise population geographic and genetic distances associated with spatial structure and serial colonization across a landscape. Likewise, IBE or “isolation-by-environment” (Wang & Summers 2010) describes the accumulation of genetic differences between environmentally distinct populations. IBE predicts that environmental differences are correlated with genetic differences, as selection differs across environments (Keller et al. 2000; Lowry et al. 2008; McBride & Singer 2010), providing a proxy for the relationship between genetic and environmental distance (Dobzhansky 1937; Wang & Bradburd 2014). The influence of geographic and environmental variation in structuring patterns of genetic variation, either independently or collectively, has received extensive support across taxa (summarize in Sexton et al. 2014). Given these observations, spatial and environmental data may provide valuable proxies in designing *ex situ* conservation collections that optimize the preservation of neutral and nonneutral evolutionary processes.

The impact of IBD and IBE on population genetic structure is expected to differ for neutral and adaptive genetic variation (Table 1). This includes the prediction that IBD will have a greater influence at neutral loci relative to IBE. IBD reflects past and current demographic history, as well as the interplay between drift and gene flow in structuring genetic variation, whereas IBE is influenced by natural selection, largely reflecting adaptive genetic variation. Cumulatively, we predict that IBD and IBE will explain the greatest proportion of genetic differences among populations for nonneutral loci. Finally, for those genetic markers underlying functional genetic diversity, including polymorphisms within genes or expressed sequences, we predict patterns of IBE and IBD will be intermediate as they may reflect a combination of adaptive and neutrally evolving loci.

**Table 1.** Evolutionary processes ^a^ contributing to genetic structure across neutral and adaptive genetic markers and their predicted weight ^b^ on expected patterns of among-population genetic differentiation (Random, IBD and IBE).

| **Neutral genetic markers** | Random | IBD | IBE |
| --- | --- | --- | --- |
| Stochastic processes (e.g. genetic drift, inbreeding) | ++ | - | - |
| Demographic history (e.g. founder events) | ++ | + | - |
| Genetic drift combined with gene flow | - | +++ | - |
| Natural selection | - | - | + |
| **Adaptive genetic markers** | Random | IBD | IBE |
| Stochastic processes (e.g. genetic drift, inbreeding) | - (+) | - | - |
| Demographic history (e.g. founder events) | - | - | - |
| Genetic drift combined with gene flow | - | + | - |
| Natural selection | - | - | +++ |

<sup>a</sup> Here we distinguish between genetic drift alone as a stochastic evolutionary force and genetic drift combined with gene flow as a process leading to a pattern of IBD.
<sup>b</sup> -: no, +: small, ++: intermediate and +++: important influence of the evolutionary forces on the pattern.

The explosion of genetic and genomic datasets publicly available provides a timely opportunity to compare the contribution of IBD and IBE to genetic structure. In the present study, we compare the influence of genetic marker type on IBD and IBE. We classify single-sequence repeats (SSRs) and genome-wide single-nucleotide polymorphisms (SNPs) as neutral genetic variation (neutral class), SNPs identified previously as candidate loci for selection using statistical or empirical methods as underlying adaptive genetic diversity (adaptive class), and genetic markers within known genes or expressed sequences (genic SNPs or expressed sequence tag SSRs) as a functional class. We distinguish functional polymorphisms from neutral and adaptive classes as these markers estimate quantitative genetic variation and likely represent a combination of neutral and adaptive processes.

To optimize sampling of genetic variation and differentiation *ex situ*, we have re-analyzed existing genetic and genomic datasets to (i) quantify the impact of IBD and IBE have on population genetic structure across neutral, functional and putatively adaptive genetic datasets, and (ii) to evaluate whether inclusion of IBD and IBE during population sampling influences genetic diversity captured at neutral, functional, and adaptive loci using simulated *ex situ* collections. We use variation partitioning to disentangle the effect of IBD, IBE, their intersection, and union on population genetic structure and then simulate *ex situ* collections using geographic and environmental distance metrics to optimize genetic variation and differentiation conserved. This study advances our understanding of the role non-genetic factors play in the distribution of genetic variation across natural populations, providing new parameters to optimize *ex situ* sampling designs where genomic data may be limited or non-existent.

## Methods

### Source of genetic and geographic data

We searched the Dryad Digital Repository (https://datadryad.org/) to identify genetic or genomics datasets for plant species using three discrete search categories: “Population structure plant”, “SSR population structure” and “SNP population structure”. Following this, for inclusion in our study, a dataset or a subset of a dataset had to meet the following criteria:

1. Populations were collected range-wide or were sampled across an isolated fraction of a species’ distribution.
2. Geographic coordinates (latitude, longitude) were available for each population sampled.
3. Genetic data, categorized as SSRs (single-sequence repeats), EST-SSRs (expressed sequence tag SSRs) or SNPs (single-nucleotide polymorphism), were available.

Range-wide sampling or sampling of populations spanning a large isolated fraction of a species’ distribution were required to ensure the majority of a species’ ecological niche space was captured. In addition, sampling a broad range of environmental and geographic distances can reduce the likelihood of covariance between environmental and geographic factors (Wang & Bradburd 2014). Using publicly available databases, population-specific latitude and longitude were used to model climatic variation associated with geographic provenance. These data were used in variation partitioning analyses and to calculate pairwise population environmental and geographic distances for each species. To calculate genetic distances, we included studies using SSRs, SNPs or EST-SSRs. SNP genotyping varied across studies, therefore we divided SNP datasets into two categories: SNPs assessed genome-wide (SNPs) and SNPs assessed within genes (Gen-SNPs). If specific SNPs were identified as being under selection based on previous work, we included a fifth category, SEL-SNPs. Finally, genetic markers were broadly classified as either putatively neutral (neutral class: SSRs, SNPs), underlying functional variation (functional class: EST-SSRs, Gen-SNPs) or putatively adaptive (adaptive class: SEL-SNPs).

Overall, we gathered 17 genetic or genomic datasets, in addition to two genomic datasets received directly from Holliday et al. (2010) (Table 2; Appendix S1). To meet the above criteria, datasets associated with seven of the 15 studied species were sub-sampled and individual geographic coordinates for one study were averaged to create population-scale coordinates (Table 2; Appendix S2).

**Table 2.**
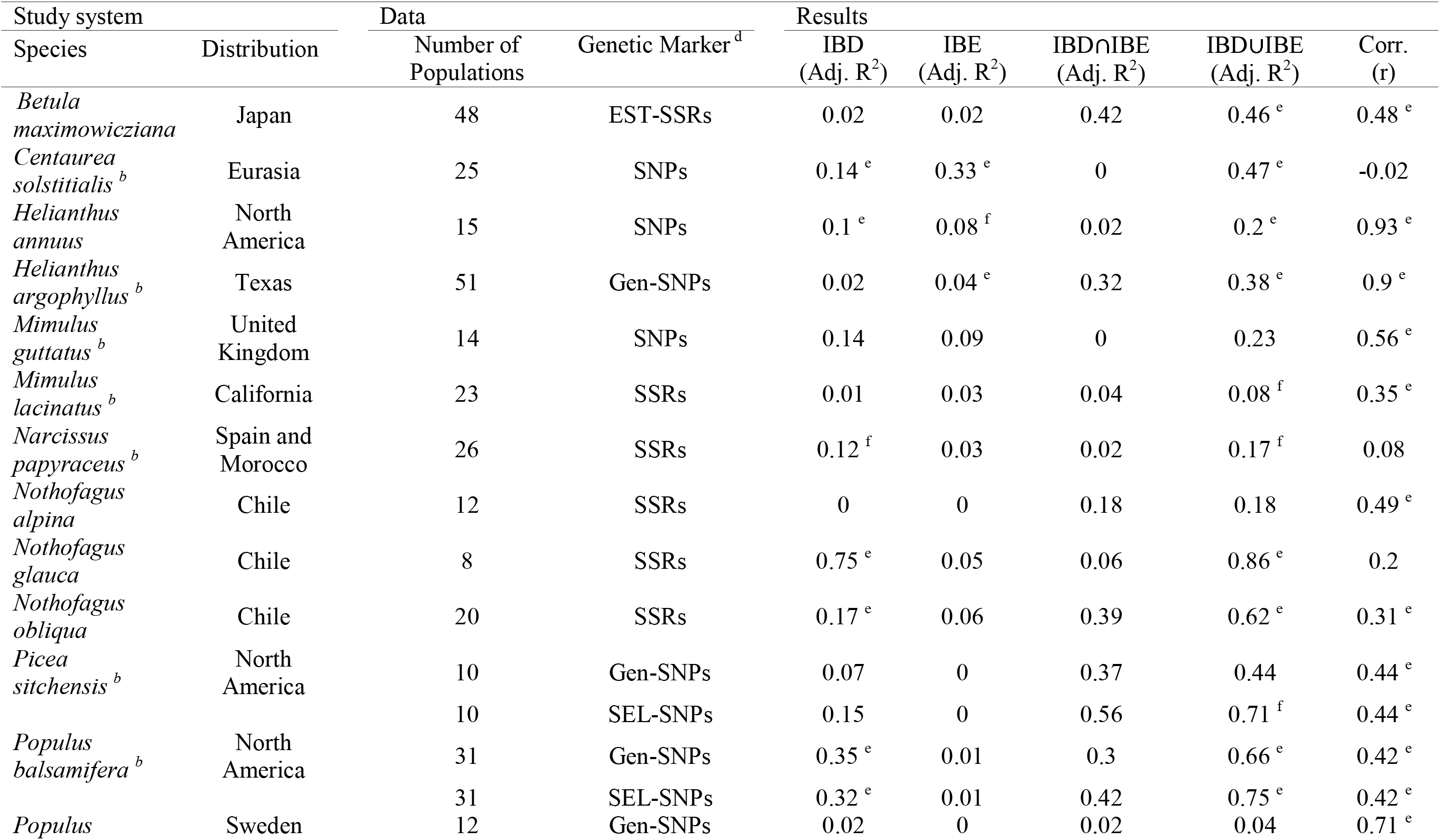
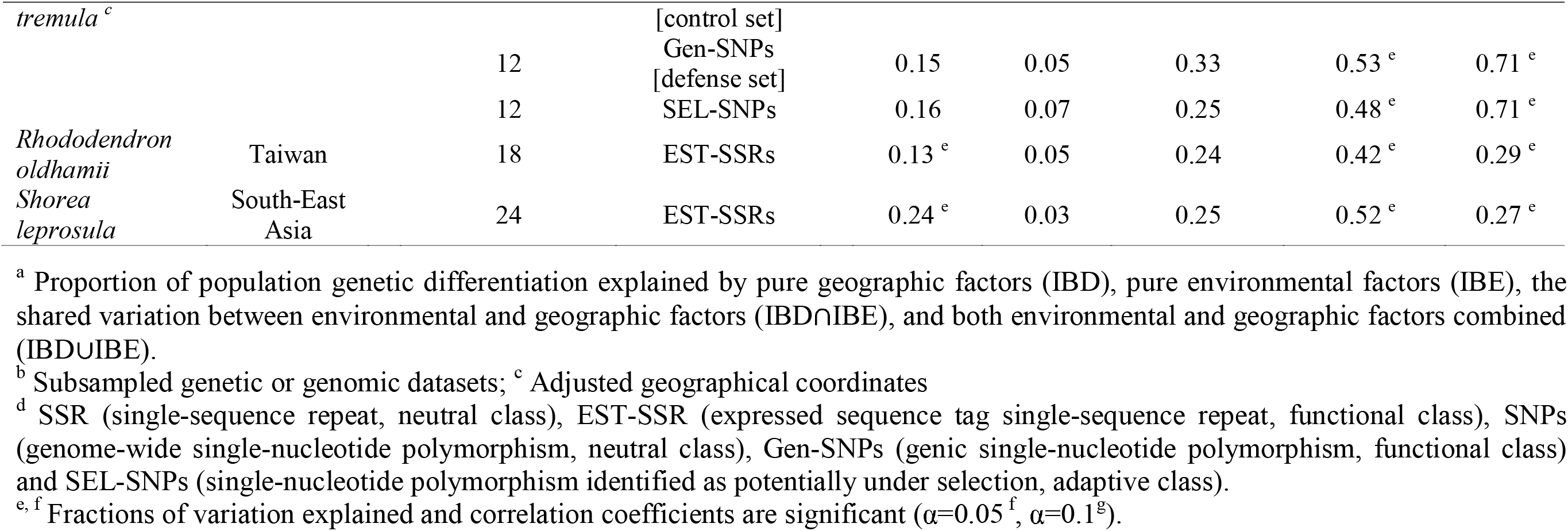
Proportion of genetic differentiation explained by environmental and geographic variables ^a^, obtained using variation partitioning analyses, and correlation coefficients estimated between pairwise geographic and environmental Euclidean distances for all 19 genetic and genomic datasets downloaded from Dryad (see Appendix S1).

### Environmental data

We used latitude, longitude and elevation associated with population provenance to extract annual, seasonal, and monthly climate variables using ClimateNA (North America), ClimateSA (South America), ClimateEU (Europe) or ClimateAP (Asia Pacific) (https://sites.ualberta.ca/~ahamann/data.html) (Appendix S3). Where elevation was not provided, GPS Visualizer (http://www.gpsvisualizer.com/elevation) was used to assign population elevation values. In total, 80 environmental variables were assigned to each population; including 79 climate-related variables and elevation. For each of the species, all environmental variables associated with population origin were filtered, standardized, and transformed to summarize environmental differences among populations. First, dataset-specific environmental variables exhibiting no population-level variation were excluded from analyses. Environmental variables were then standardized and used to conduct a principal component analysis (PCA). PCA was used to reduce the overall number of environmental variables by summarizing environmental differences across two major axes of differentiation, which together explain more than 70% of environmental variation observed between populations (Appendix S4). These two major PC axes were considered as predictor variables for variation partitioning and used to calculate population pairwise environmental distances in simulations.

### Variation partitioning analysis

To quantify the contribution of IBD and IBE to genetic divergence within each of the 19 datasets, we conducted a variation partitioning analysis in R (R core Team 2018) using the “vegan” package (Oksanen et al. 2007). We used standard estimates of population genetic differentiation re-calculated for all population pairs within each dataset as our response variable. To account for variation in genetic markers, we used Nei’s F_ST_ (Nei 1987), as this metric can provide comparable estimates of population genetic differentiation for both biallelic (e.g. SNPs) and multi-allelic (e.g. SSRs) loci. For each dataset, population divergence was partitioned between two sets of predictor variables; including the geographic coordinates (latitude, longitude) and the two major environmental PC axes (PC1, PC2) associated with each population within a dataset. Following variation partitioning, we conducted a partial distance-based redundancy analysis (dbrda) on each dataset to test the significance of (i) variance explained by each set of predictor variables alone (IBD, IBE; Table 2), and (ii) the variance explained by the union of predictor variables (IBD⋃IBE; Table 2). We did not evaluate the significance of the variance explained by the intersection of geographically structured environmental variables (IBD⋂IBE; Table 2), as this variance fraction is not testable using dbrda.

### Quantifying the correlation between genetic, environmental and geographic distances

Geographic and environmental distance between population pairs was measured as the Euclidean distance between populations’ geographic coordinates (latitude, longitude) or. between populations’ two major environmental PC axes (PC1, PC2), respectively. To visualize and evaluate the covariance structure between genetic, environmental and geographic distance matrices, we graphed and estimated the correlation between all distance metrics (Table 2; Appendix S5). Correlation coefficients were estimated using the nonparametric mantel test implemented in the R package “adegenet” (Jombart 2008) for each dataset separately.

### Simulating an *ex situ* collection: an idealized framework

We simulated an idealized *ex situ* conservation collection for each dataset using a customized R script relying on R packages “adegenet” (Jombart 2008), “hierfstat” (Goudet 2005) and “data.table” (Dowle & Srinivasan 2019). This simulation measured the amount of genetic differentiation and the proportion of allelic diversity captured in *ex situ* collections that prioritize population sampling based on environmental and geographic distances. We simulated *ex situ* collections using four different population sampling strategies. This included random sampling, as well as sampling prioritized based on distances between populations’ two major environmental PC axes (Euclidean environmental distance), sampling based on distances between populations’ geographic coordinates (Euclidean geographic distance) or both (Fig. 1a).

**Figure 1.**
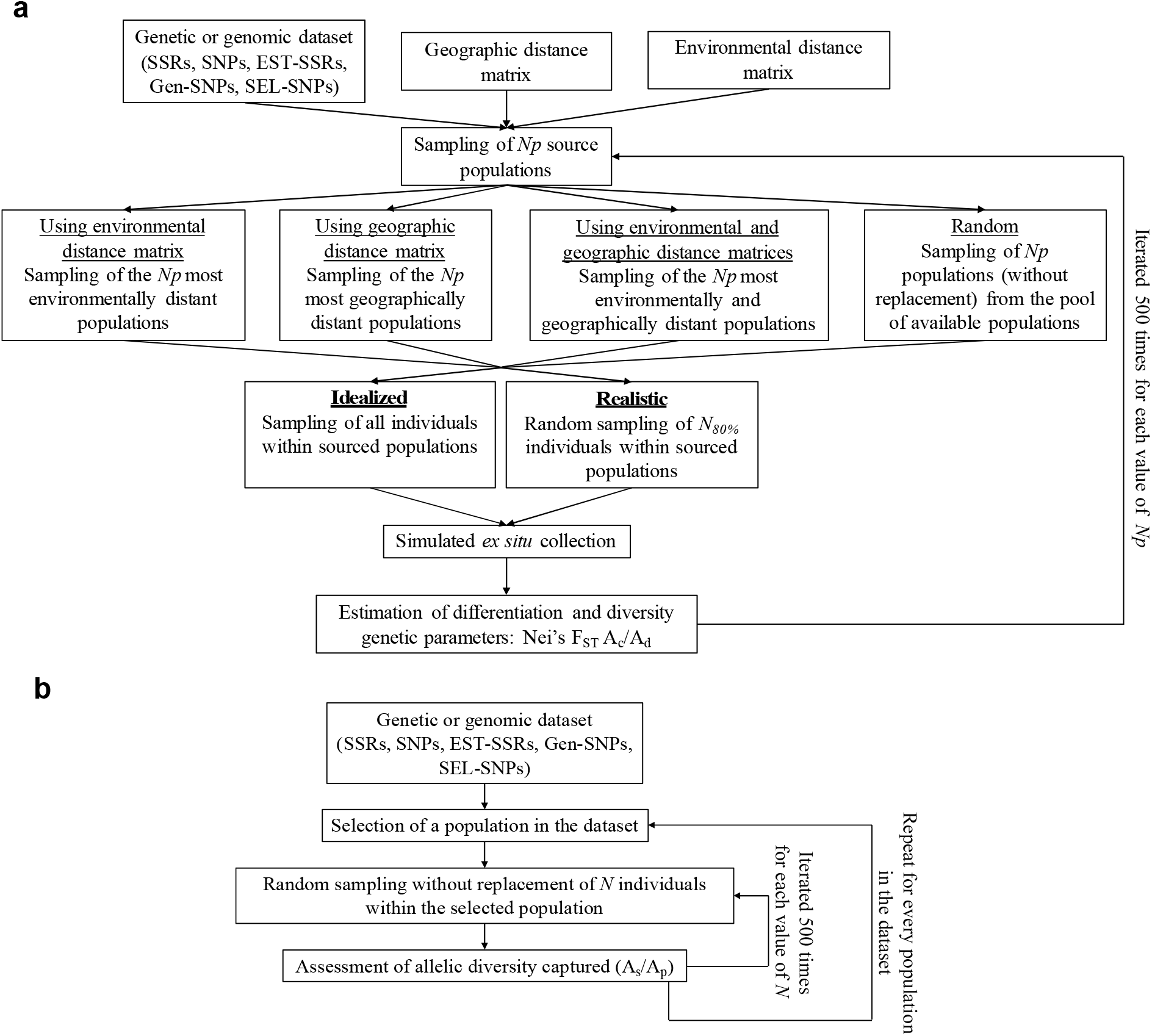
(a) Simulation framework used to estimate genetic variation and differentiation parameters in *ex situ* collections simulated under two different within-population sampling scenarios (realistic and idealized) and four distinct population prioritization strategies (random, based on environmental distance, based on geographic distance, and based on both environmental and geographic distance combined). (b) Simulation framework used to estimate the number of individuals required to capture between 80-100% of allelic diversity in every population of a dataset (*N*_*80%*_, see Figure 1a). Simulations using both frameworks were conducted on each dataset independently. Computation proceeds from top to bottom.

*Ex situ* collections were simulated using between two and the total number of populations available for each dataset (*Np*, Fig. 1a). Randomized sampling sampled populations without replacement from the pool of available populations. Environmentally or geographically prioritized simulations sampled population pairs with the greatest pairwise distances in decreasing order. Collections simulated using the combination of environmental and geographic distances sampled population pairs that exhibited the greatest sum of environmental and geographic distances following standardization, prioritized in decreasing order. All individuals within each population were sampled as part of the idealized simulation.

To compare genetic diversity captured across simulated collections, we estimated two genetic parameters: Nei’s F_ST_ and allelic diversity captured (A_c_/A_d_). These indices were chosen as they quantify different aspects of population genetic diversity. Nei’s F_ST_ provides an estimate of genetic differentiation across sampled populations and A_c_/A_d_ provides an estimate of the number of alleles captured in collections (A_c_) relative to the total number of alleles present within a dataset (A_d_). All genetic parameters were estimated in R using the “hierfstat” package.

Population sampling and associated genetic summary statistics were simulated 500 times for each dataset to account for the variance introduced through randomly sampling across populations. Summary statistics were estimated based on average values across all 500 simulations. No replication was used for environmental and/or geographic distance-based population sampling, as neither provenance of source populations nor genetic summary statistics would have changed with repeated iterations.

For these idealized simulations, all individuals were sampled within each target population (equivalent to protecting the entire population), regardless of collection strategy, assuming 100% of the standing genetic variation was captured. However, monetary or logistical constraints usually impact the number of individuals that could be sampled within a target population. Given this, we predict that genetic diversity captured within source populations will vary. To assess whether insights gained from idealized simulations were maintained under more realistic conditions, we conducted additional simulations, introducing differences in the amount of genetic diversity captured between populations (hereafter referred to as realistic simulations).

### Simulating an *ex situ* collection: The realistic framework

To simulate a realistic *ex situ* collection, a subset of individuals was sampled within each population. This provides the opportunity to evaluate the impact varying genetic diversity captured within populations may have on total genetic diversity and differentiation captured across populations collected. We assume that *ex situ* collections aim to preserve as much genetic variation as possible within each population. Within this framework, we postulated that at least 80% of within-population allelic diversity would be captured *ex situ*. Therefore, for each dataset, we assessed the number of individuals (*N*_*80%*_) that when sampled capture between 80%-100% of allelic diversity across populations.

An additional simulation was used to determine the value of *N*_*80%*_ for each dataset (Fig. 1b). For every population, *N* individuals (ranging from one up to the size of the smallest population within the assessed dataset) were randomly sampled without replacement. Following this, the number of alleles captured for *N* individuals (A_s_) divided by the total number of alleles in the population (A_p_) was quantified for each population. Sampling of individuals and quantification of allelic diversity captured was replicated 500 times for each population and value of *N* to calculate confidence intervals around A_s_/A_p_ ratios. The number of individuals required to capture 80% or more (As/Ap ≥0.8) of allelic diversity in every population (*N*_*80%*_) was visually assessed for each dataset independently (Appendix S6) and used to parametrize realistic simulations (Fig. 1a). *Ex situ* collections were simulated 500 times using the realistic scenario to estimate genetic summary statistics regardless of the population sampling strategy used (Fig. 1a). For these simulations, *N*_*80%*_ were often much lower than the existing size of most populations and performing repeated iterations accounted for the variation in genetic summary statistics introduced by small values of *N*_*80%*_.

Maintaining the range of A_s_/A_p_ ratios across datasets is crucial as unbalanced variance may confound the influence of prioritization strategies in downstream analyses. Four of the 19 datasets (*H. argophyllus* (Gen-SNPs), *M. lacinatus* (SSRs), *R. oldhamii* (EST-SSRs) and *S. leprosula* (EST-SSRs)) were discarded from realistic simulations as *N*_*80%*_ values were not reached for these datasets (Appendix S6). These same datasets were also removed from idealized simulations to ensure that differences in summary statistics between idealized and realistic simulations originated solely from variation in allelic diversity captured across populations introduced in the latter. See Appendix S7 for a complete list of parameters tested and used for simulations.

### Analysis of simulated data

We tested whether prioritizing source population collection using environmental and/or geographic distance data influences genetic variation and differentiation captured *ex situ*. For every number of populations sampled (*Np*), genetic summary statistics simulated using random sampling were subtracted from values based on prioritization strategies using environmental distances, geographic distances, or both. Summary statistics were averaged for each dataset following repeated iterations, grouped by distance-based strategies, genetic marker class, and simulation framework (idealized or realistic) (Fig. 2). Differences in genetic summary statistics are provided based on the proportion of populations sampled as the number of populations sampled for analysis varied across studies. For each dataset, we selected four numbers of populations sampled (*Np*) representing between 30-40%, 50-60%, 70-80%, and 90-100% of populations present in a dataset (Appendix S7).

**Figure 2.**
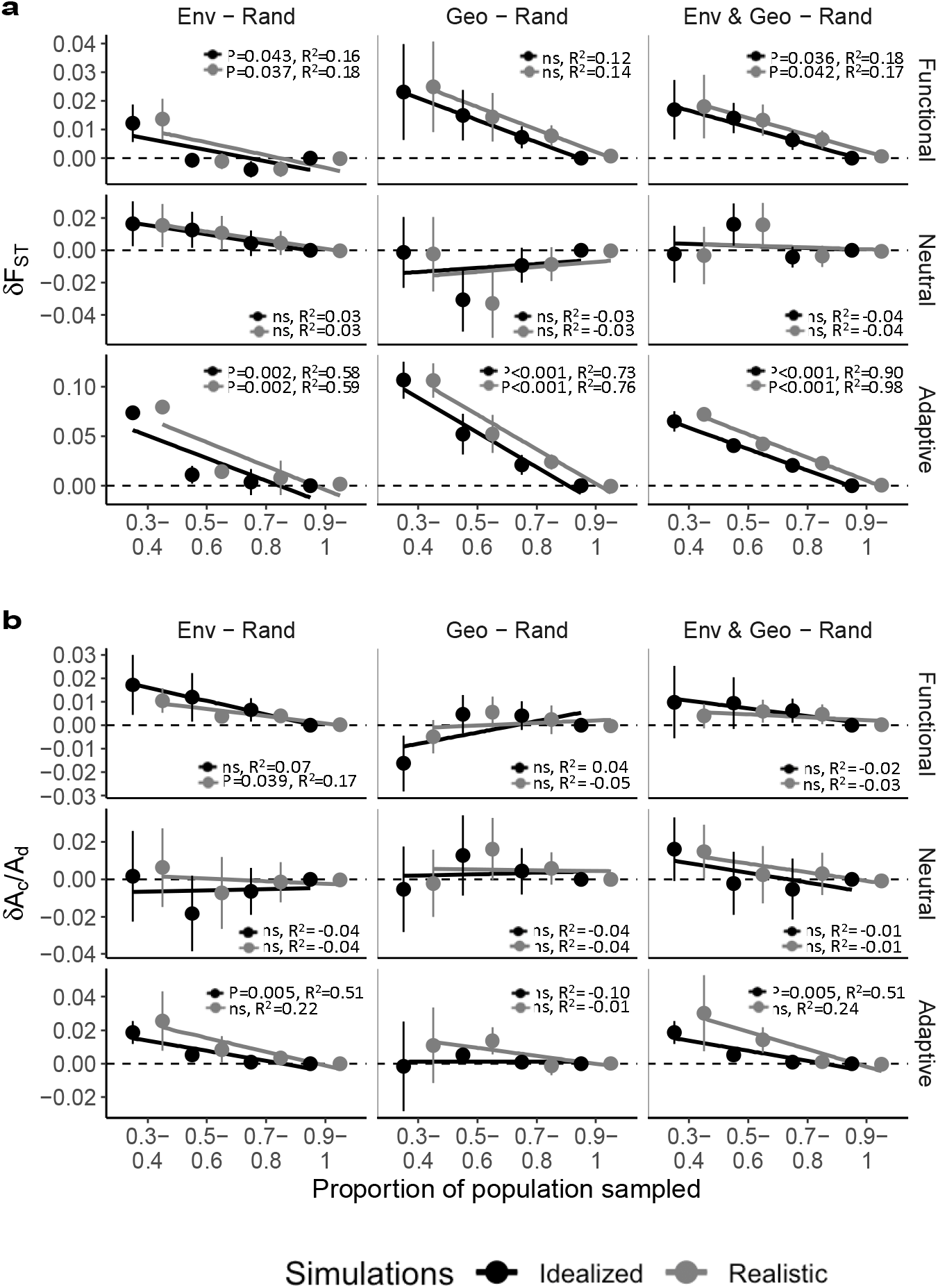
Average differences and SE across datasets in genetic summary statistics (y-axis) estimated from *ex situ* collections simulated using distance-informed (environmental: Env, geographic: Geo, environmental and geographic: Env & Geo) and random (Rand) population sampling strategies (columns) separated by genetic marker classes (rows). Differences in genetic summary statistics were estimated for various proportions of populations sampled (x-axis). (a) Populations genetic differentiation (Nei’s F_ST_). (b) Allelic diversity captured in simulated *ex situ* collections (A_c_/A_d_). ns: non-significant.

Finally, we fitted a linear model between proportions of populations sampled and differences in genetic summary statistics for every combination of genetic marker class, distance-based prioritization strategy, and simulation framework (Fig. 2). A negative relationship indicates that a given distance-informed sampling generally increases the genetic summary statistics relative to random sampling while a positive relationship would suggest the opposite. In addition, it is important to note that a significant relationship (positive or negative) will always be approaching zero as the proportion of populations sampled increases. This is because with additional populations sourced, the probability that identical populations are sampled randomly or via distance-based strategies increases and will reach one when all populations are sampled. As the number of shared populations between sampling strategies increases, the difference in genetic summary statistics decreases.

## Results

### Relative contributions of IBD and IBE to population genetic differentiation

Variation partitioning revealed that IBD explained significantly more among-population genetic differences (13%) than IBE alone (5.5%) or IBD⋂IBE (3%) for neutral genetic datasets (Table 3). This contrasts with functional and adaptive datasets, where a significant proportion of among-population genetic differences was explained by geographically structured environmental variables relative to environmental or geographic factors alone (Table 3). Overall, 31% and 42% of population genetic differences were explained by IBD⋂IBE for functional and adaptive datasets, respectively, while only a small proportion was explained by IBD (functional: 10%, adaptive: 16%) and IBE alone (functional: 2.5%, adaptive:1%).

**Table 3.** Median proportion and 95% CI* of population genetic differences explained by IBD, IBE, IBD⋂IBE, and IBD⋃IBE given by genetic marker classes.

| Genetic Marker Class | IBD | IBE | IBD∩IBE | IBD∪IBE |
| --- | --- | --- | --- | --- |
|  | Median Adj. R<sup>2</sup> (95% CI) | Median Adj. R<sup>2</sup> (95% CI) | Median Adj. R<sup>2</sup> (95% CI) | Median Adj. R<sup>2</sup> (95% CI) |
| Neutral | 0.13 (0.09, 0.25) | 0.055 (0.02, 0.08) | 0.03 (-0.12, 0.06) | 0.215 (-0.19, 0.26) |
| Functional | 0.1 (-0.04, 0.18) | 0.025 (0, 0.045) | 0.31 (0.25, 0.38) | 0.45 (0.37, 0.52) |
| Adaptive | 0.16 (0, 0.17) | 0.01 (-0.05, 0.02) | 0.42 (0.28, 0.59) | 0.71 (0.67, 0.94) |

<sup>\*</sup> 95% CI were obtained by bootstrapping. We considered two medians to be significantly different ( $\alpha=0.05$ ) if their confidence intervals did not overlap.

While significant differences in the proportion of genetic differentiation explained were observed across genetic marker classes for IBD⋂IBE and IBD⋃IBE, no significant differences were observed in the individual contribution of IBD and IBE (Table 3). IBD⋃IBE explained the greatest proportion of genetic differences for adaptive genetic markers (71%), followed by functional (45%) and neutral (21.5%) genetic markers, respectively. Interestingly, IBD⋂IBE explained substantial among-population genetic differences for both functional and adaptive datasets but explained limited variation for neutral datasets (Table 3). The contribution of IBD⋂IBE to population genetic differentiation for adaptive and functional datasets likely reflect high correlations observed between environmental and geographic distance matrices (Table 2; Appendix S5). Therefore, the relative contribution of geography and environment should be interpreted with caution for these genetic marker classes, as population genetic differentiation could not be partitioned solely by IBD or IBE.

### Genetic diversity and differentiation captured in simulated *ex situ* collections

#### Genetic differentiation (Nei’s F_ST_)

Significant negative relationships were observed between proportions of populations sampled and changes in genetic differences (F_ST_) captured for collections simulated using both adaptive and functional datasets, but not neutral genetic datasets (Fig. 2a). This suggests that using environmental and/or geographic distance to prioritize population sampling may potentially increase adaptive and functional genetic differences but does not consistently impact neutral genetic variation. Simulations revealed that using all three distance-based population sampling strategies increased genetic differentiation captured among adaptive loci in *ex situ* collections (Fig. 2a). This contrasts with the results obtained for functional datasets, where sampling prioritizing source populations using environmental distance, or the combination of both environmental and geographic distances increased genetic differences captured.

For both adaptive and functional genetic makers classes, simulations based on realistic and idealized within-population sampling scenarios led to similar slopes, regardless of the distance-based population sampling strategy used (Fig. 2a; Appendix S8). This indicates that the ability of distance-based population sampling strategies to increase F_ST_ among functional and adaptive loci was not impacted by the within-population sampling scenarios simulated.

#### Proportion of allelic diversity captured (A_c_/A_d_)

Both realistic and idealized *ex situ* collection simulations using functional and adaptive genetic datasets indicated allelic diversity captured (A_c_/A_d_) is likely sensitive to within-population sampling. Prioritizing population sampling using environmental distances increased allelic diversity captured at functional loci under realistic within-population sampling conditions, but had no impact using idealized within-population sampling scenario (Fig. 2b). This contrasts with results obtained for adaptive datasets, where the opposite pattern was observed. Prioritizing population sampling using environmental or the combination of environmental and geographic distances increased A_c_/A_d_ under idealized within-population sampling conditions (Fig. 2b).

For neutral genetic datasets no consistent change in allelic diversity was observed in response to varying proportions of population sampled, regardless of population prioritization strategy tested or within-population sampling scenario simulated (Fig. 2b). Together, these results suggest that incorporating environmental and/or geographic distances to prioritize collections may increase allelic diversity captured at functional and adaptive loci, but not at neutral loci. Nonetheless, simulations also indicate that increasing allelic diversity captured in *ex situ* collections is dependent on within-population sampling scenarios and may thus only be achieved under specific sampling conditions.

## Discussion

Optimizing efforts to conserve genetic variation relies upon an understanding for how non-genetic factors, geographic and environmental variation, contribute to population genetic structure. Here, we leverage population provenance and environmental data to optimize genetic differences captured in simulated conservation collections. Environmental and geographic factors explain some portion of the genetic differences observed among populations, although the extent differs by genetic marker class. The proportion of genetic differentiation explained by IBD⋃IBE was significantly higher for adaptive and functional datasets relative to neutral datasets. This suggests that geographic and environmental data may provide a useful guide when designing *ex situ* population sampling, particularly where the goal is to conserve adaptive and functional genetic variation. We simulated *ex situ* sampling and found that, as predicted, strategies that included environmental and/or geographic distance data to prioritize population sampling increased genetic differences and diversity captured at both functional and adaptive loci. Overall, we suggest that inclusion of IBD and IBE in guiding *ex situ* sampling can ensure adaptive and functional genetic variation are conserved, crucial for long-term preservation and maintenance of species’ evolutionary potential.

Consistent with previous plant studies, our results demonstrate that genetic differentiation across neutral, functional, and adaptive loci can, at least partly, be explained by environmental and geographic factors (Bjørnstad et al. 1995; Nadeau et al. 2016; Xia et al. 2018) (Table 2). Interestingly, limited genetic differentiation was explained by IBD or IBE alone across all three genetic marker classes. For functional and adaptive datasets, this is likely due to the fact that substantial genetic structure is explained by their intersection (Table 3). Indeed, IBD⋂IBE reflects covariance between geographic and environmental factors that cannot be teased apart. Additional empirical work minimizing this covariance would be required to completely disentangle these factors (Wang & Bradburd 2014). Nonetheless, when combined, environmental and geographic factors explained a substantial proportion of population genetic differentiation for both functional and adaptive datasets (IBD⋃IBE; Table 3). This suggests that geographic and environmental differences contribute largely to genetic divergence at nonneutral loci (Huang et al. 2016; Xia et al. 2018). Consequently, the inclusion of IBD⋃IBE may provide a means to capture adaptive and functional genetic variation *ex situ*. For neutral datasets, geographic and environmental factors, either individually (IBD, IBD) or cumulatively (IBD⋃IBE), explained very small proportions of among-population genetic differences (Table 3). This indicates that stochastic processes, such as genetic drift or founding events likely influence neutral genetic structure. Random fixation or loss of alleles through genetic drift (Stern & Orgogozo 2009) and accelerated allele fixation within populations following demographic changes, including bottlenecks or founder events (Maruyama & Fuerst 1985; Gavrilets & Hastings 1996), may lead to population structure that is not explained by environment or spatial data. Overall, our findings indicate that environmental and geographic distance metrics can be used to target genetic differences which likely reflect adaptive or functional genetic variation over neutral genetic variation.

*Ex situ* strategies relying on existing genetic datasets (Caujapé-Castells & Pedrola-Monfort 2004; Gapare et al. 2008) or genetic simulations (Hoban & Schlarbaum 2014; Hoban 2019) have previously optimized variation captured in collections. These approaches require substantial *a priori* information and target neutral genetic variation. Where knowledge of population location is available, pairwise geographic and environmental distances may be leveraged to extend previous sampling to conserve adaptive and functional genetic variation. Our simulations demonstrate that *ex situ* collections prioritized using environmental or the combination of environmental and geographic distances increase both Nei’s F_ST_ and A_c_/A_d_ captured for adaptive and functional datasets relative to random sampling (Fig. 2). This indicates that divergent selection and adaptation to local environments contribute to genetic differentiation at nonneutral loci (Hancock et al. 2011; Wang et al. 2016), likely influenced by IBE. IBE-based prioritization strategies suggest that part of the additional genetic differences captured in collections consist of spatially and/or environmentally restricted alleles (Fig. 2b). However, simulations also revealed that increasing allelic diversity captured in collections using distance-based prioritization strategies depends on within-population sampling conditions (realistic or idealized). These results have important applications to applied conservation efforts. First, a realistic sampling scenario was sufficient to increase genetic differentiation captured at adaptive and functional loci (Fig. 2a). This suggests that inclusion of IBD and IBE in population prioritization would likely increase among-population genetic differences captured at these loci by sampling only a subset of individuals within populations. However, only an idealized sampling scenario increased allelic diversity captured at adaptive loci (Fig. 2b). This indicates that extensive within-population sampling may be needed to increase adaptive allelic diversity conserved in collections. Overall, simulations demonstrate that prioritizing population sampling using IBD and/or IBE can increase genetic differences and diversity captured at both functional and adaptive loci without the need for prior genetic data, providing a means to target genetic variation that may be needed to maintain adaptive potential within collections.

Despite the fact conservation has long valued environmental and geographic data (Brown & Marshall 1995; Guerrant et al. 2004; Guerrant Jr et al. 2013), use of these data for conservation planning have only emerged during the past decade (Vinceti et al. 2013; Whitlock et al. 2016; Hanson et al. 2017). Consistent with previous work, we observe inconsistent benefits of leveraging geography for the preservation of neutral genetic diversity (Fig. 2). This could be due to the fact that gene flow between populations may be disturbed by landscape characteristics (Dudaniec et al. 2016), or some species may exhibit greater gene flow between geographically distant populations (O’Connell et al. 2007). Our results do provide additional empirical support for inclusion of environmental and geographic data in conservation planning, to target and increase adaptive genetic diversity conserved (Hanson et al. 2017) (Fig. 2). In addition, this study is the first to provide evidence that IBD- and/or IBE-based population prioritization strategies may increase genetic differentiation and diversity captured at functional loci. This indicates that using environmental and/or geographic surrogates may not only preserve current adaptive genetic diversity but may also secure genetic variation crucial for future adaptations. Finally, where other studies use amplified fragment length polymorphisms (AFLPs; Whitlock et al. 2016; Hanson et al. 2017), we focus on SSRs and SNPs datasets. The concordance across studies suggests a broad applicability for environmental and geographic data to act as surrogates to optimize the conservation of genetic variation.

Although simulations are a powerful inferential tool, they can include a number of assumptions. Here, we assumed that maternal plants used in realistic and idealized simulations were collected for storage *ex situ*. However, the progeny of these plants more accurately reflects those likely to be included in collections (FAO, 2010). Future studies will need to consider empirical or simulated progeny data to evaluate whether environmental and/or geographic distance-based prioritization captures genetic variation across generations. In this study, we evaluated the overall impact of population sampling strategies on genetic variation and differentiation captured in *ex situ* collections. Nonetheless, simulations revealed important variation in genetic summary statistics across datasets within genetic marker classes (Fig. 2). This variation is likely introduced by differences in species’ life history traits including mode of reproduction and breeding system (Loveless & Hamrick 1984). Despite this variance, our data suggest that inclusion of IBD and IBE in *ex situ* guidelines may still be valuable to optimizing functional and adaptive genetic variation captured. Future work assessing the influence trait combinations may have on predicting genetic variation captured in collections will complement the present research, providing sampling guidelines for species exhibiting specific life history characteristics. Finally, we grouped different genetic markers into genetic diversity classes to test the effect of prioritizing population sampling using environmental and/or geographic data at a broader scale. However, allelic distributions and mutation models largely differ between these genetic markers. Thus, future work should evaluate marker-specific patterns associated with IBD- and IBE-based prioritization strategies.

Anthropogenic changes have had substantial impacts on global biodiversity, resulting in a global call for the preservation of biodiversity. This research expands existing *ex situ* population sampling strategies, leveraging geographic provenance and environmental distance to increase functional and adaptive genetic differences conserved in collections. Incorporating an understanding of evolutionary and ecological processes influencing population structure alongside new and existing datasets will be critical to enhancing current conservation practice.

## Supporting Information

Reference and availability information associated with every genetic and genomic dataset (Appendix S1), modifications applied to genetic and genomic datasets (Appendix S2), raw set of climatic variables used in simulations and variation partitioning analyses (Appendix S3), proportion of variance explained by the two major environmental principal components for each dataset (Appendix S4), covariance between environmental, geographic and genetic distances (Appendix S5), proportion of allelic diversity captured within populations using *N*_*80%*_ or the size of the smallest population within datasets (Appendix S6), a list of tested and used parameters for realistic and idealized simulations (Appendix S7), and regression statistics associated with realistic and idealized simulations (Appendix S8) are available online.

## Acknowledgments

The authors thank the Hamilton Lab, T. L. Parchman, and S. M. Hoban for their valuable comments on early versions of this manuscript. This work was supported by a new faculty award from the office of the North Dakota Experimental Program to Stimulate Competitive Research (ND-EPSCoR NSF-IIA-1355466) and funding from the NDSU Environmental and Conservation Sciences Program to J.A.H.

