## Supplementary Information S1-S8 for "Environmental and geographic data optimize *ex situ* collections and the preservation of adaptive evolutionary potential"

### Appendix S1 – Reference and availability information associated with genetic and genomic datasets of all 15 plant species under study.

| Species | Literature reference | Dryad link |
| --- | --- | --- |
| *Betula maximowicziana* | (Tsuda et al. 2015) | <https://doi.org/10.5061/dryad.dj17c> |
| *Centaurea solstitialis* | (Barker et al. 2017) | <https://doi.org/10.5061/dryad.pf550> |
| *Helianthus annuus* | (McAssey et al. 2016) | <https://doi.org/10.5061/dryad.6p1c4> |
| *Helianthus argophyllus* | (Moyers & Rieseberg 2016) | <https://doi.org/10.5061/dryad.3c769> |
| *Mimulus guttatus* | (Pantoja et al. 2017) | <https://doi.org/10.5061/dryad.91v3n> |
| *Mimulus lacinatus* | (Sexton et al. 2016) | <https://doi.org/10.5061/dryad.8qc40> |
| *Narcissus papyraceus* | (Simón‐Porcar et al. 2015) | <https://doi.org/10.5061/dryad.jh21r> |
| *Nothofagus alpina* | (Vergara et al. 2014) | <https://doi.org/10.5061/dryad.h3d26> |
| *Nothofagus glauca* | (Vergara et al. 2014) | <https://doi.org/10.5061/dryad.h3d26> |
| *Nothofagus obliqua* | (Vergara et al. 2014) | <https://doi.org/10.5061/dryad.h3d26> |
| *Picea sitchensis* | (Holliday et al. 2010) | * |
| *Populus balsamifera* | (Keller et al. 2017) | <https://doi.org/10.5061/dryad.gp78p> |
| *Populus tremula* | (Bernhardsson & Ingvarsson 2012) | <https://doi.org/10.5061/dryad.0vr6m66t> |
| *Rhododendron oldhamii* | (Hsieh et al. 2013) | <https://doi.org/10.5061/dryad.nc221> |
| *Shorea leprosula* | (Ohtani et al. 2013) | <https://doi.org/10.5061/dryad.1mt1h> |

* Datasets received directly from the authors.

### Appendix S2 – Modifications applied to species datasets.

Genetic and genomics datasets associated with 7 of the 15 studied species downloaded from Dryad (<https://datadryad.org/>) were modified to meet selection criteria for subsequent analysis. Modifications performed involved subsampling of original datasets through the removal of one or several individuals, populations or genetic markers. In addition, geographic coordinates provided in Bernhardsson & Ingvarsson (2012) also needed to be adjusted to fully meet the second selection criterion. Below is a detailed list of modifications made to genetic, genomic, and geographic datasets.

*Centaurea solstitialis –* First, individuals belonging to another species that *C. solsitialis* (*Centaurea melitensis*, *Centaurea nicaeensis,* and *Centaurea pallescens*) were discarded because they did not span species geographic ranges or isolated part of their ranges and therefore violated the first selection criterion. Second, only the individuals of *C. solstitialis* distributed in Eurasia were kept because samples from the United States and South America were not representative of the species distribution or an isolated part of its distribution in these locations, also violating the first selection criteria. Third, individuals with the prefix “Ar-” were discarded because they could not be associated with a unique population. Two populations in Barker et al. (2017) were named “AR”, on located in the United States (Lat 45.696˚, Long −118.871˚), the other located in South America (Lat 39.563˚, Long 67.007˚). Additionally, individuals labeled C1346, C1441, C1445, and C1448 as well as individuals with the prefix “Sie-” were removed from the dataset because they could not be assigned to any populations. Finally, individuals belonging to populations ETC and K113 as well as individuals labeled as C1309, C1310, and C1313, all three belonging to the same population, were discarded as they fell without the geographic range covered by ClimateEU and could not be associated with climatic variables. This would prevent variation partitioning to be performed and environmental distance among populations to be estimated, impeding downstream statistical analyses. Overall, 225 individuals from 25 populations spanning the species native (Eurasian) distribution range met all three selection criteria and were used for analyses.

*Helianthus argophyllus –* First, all individuals from another species than *H. argophyllus* (*Helianthus annuus* and *Helianthus debilis*) were discarded as they were poorly sampled and covered only a small part of species distribution ranges, violating the first selection criterion. Second, individuals within populations labeled as ARG-1575, ARG-2623, HEL153, PI649866, PI490291_Moz, 448, 449 and 451 were removed from the dataset because no geographic coordinates were available for these populations, violating the second selection criterion. Overall, 554 individuals from 51 populations spanning the species native ancestral distribution range (Texas; Yatabe et al. 2007), which currently represents an isolated fraction of its distribution (Texas, Florida and North Carolina; [https://plants.usda.gov](https://plants.usda.gov/)) met all three selection criteria and were used for analyses.

*Mimulus guttatus* – Individuals belonging to populations labeled as ALA, WLB, CPB, HOC, HEC, ANR, LMC, WTB, DFAL and DAV were discarded as they were not representative of the species North American distribution nor representative of a fraction of the species range, violating the first selection criterion. Overall, 261 individuals from 14 populations sampled across the species British distribution met all three selection criteria and were used for subsequent analyses.

*Mimulus lacinatus –* In this dataset, 3 of 11 codominant genetic markers were removed (e617, e641, e423) as they are gene-intron-length markers and did not fall within the range of genetic markers under study (SSR, EST-SSR, SNP, Gen-SNP, SEL-SNP), violating the third selection criterion. The remaining eight microsatellite markers were used for analyses.

*Narcissus papyraceus –* Individuals within 5 of the 31 populations studied in Simón‐Porcar et al. (2015) (populations 1-5, region code CM) were discarded because they fell without the geographic range covered by ClimateEU. Consequently, no climatic variable could be retrieved for these sites, which would prevent variation partitioning to be performed and environmental distance among populations to be estimated, impeding downstream statistical analyses. Overall, 422 individuals from 26 populations sampled throughout most of the species European and North African distribution range met all three selection criteria and were used for analyses.

*Picea sitchensis –* First, individuals labeled with prefixes “10” and “15” were removed from the SNPs dataset as none of these individuals were assigned to one of the populations listed in Holliday et al. (2010). Second, of the 35 SNPs identified as putatively under selection in the study, only 34 were considered as SEL-SNPs because one SNP (273_98_NS) was absent from the original full genotypes’ dataset. Finally, individuals belonging to Rocky Bay and Kodiak Island populations were discarded as no climatic data could be retrieved for these two populations using ClimateNA, which would prevent variation partitioning to be performed and environmental distance among populations to be estimated, impeding downstream statistical analyses. Overall, 286 individuals from 10 populations spanning most of the species native (North American) distribution range met all three selection criteria and were used for analyses.

*Populus balsamifera –* First, individuals from another species than *P. balsamifera* (*Populus deltoids*, *Populus tremula*, *Populus tremuloides*, *Populus trichocarpa,* and *Populus angustifolia*) were discarded as only a few individuals were sampled (1-8 individuals) per species and thus neither represent whole nor part of species distribution ranges, violating the first selection criterion. Second, 7 genetic markers (ABi1D_183, CRY11_3201, GI5_5271, PHYB2_5048, GI2_10278, PIF31_1277, PIF31_2601) were removed from the genomic dataset as they represent indel and not SNP variation, violating the third selection criterion. Finally, to comply with the second selection criterion, only individuals belonging to the 31 populations studied in Keller et al. (2012) were kept as geographical coordinates could only be retrieved for these populations. Overall, the dataset we used for analyses matched the one used in Keller et al. (2012), including 443 individuals from 31 populations spanning most of *P. balsamifera* distribution range.

*Populus tremula –* In the study performed by Bernhardsson & Ingvarsson (2012), geographical coordinates are given per tree sampled. However, to fully comply with the second selection criterion and be able to both conduct variation partitioning and estimate environmental and geographical distances among populations, geographical coordinates per population are needed. To resolve this issue, we pooled all individuals belonging to the same population together and averaged values of latitude, longitude, and elevation to generate geographical coordinates and elevation data per population.

### **Appendix S3 -** Raw set of climatic variables assigned to populations of every genetic and genomic dataset using ClimateNA, SA, EU or AP (<https://sites.ualberta.ca/~ahamann/data.html>).

| Annual variables | Seasonal variables | Monthly variables |
| --- | --- | --- |
| Mean Annual Temperature (˚C)  Mean Warmest Month Temperature (˚C)  Mean Coldest Month Temperature (˚C)  Continentality (˚C)  Mean Annual Precipitation (mm)  Annual Heat-Moisture index  Degree-Days below 0˚C  Degree-Days above 5˚C  Degree-Days below 18˚C  Degree-Days above 18˚C  Number of frost-free days  Precipitation as snow (mm)  Extreme Minimum Temperature over past 30 years (˚C)  Hargreaves reference evaporation (mm)  Hargreaves climatic moisture deficit (mm) | Winter mean maximum Temperature (˚C)  Spring mean maximum Temperature (˚C)  Summer mean maximum Temperature (˚C)  Autumn mean maximum Temperature (˚C)  Winter mean minimum Temperature (˚C)  Spring mean minimum Temperature (˚C)  Summer mean minimum Temperature (˚C)  Autumn mean minimum Temperature (˚C)  Winter mean Temperature (˚C)  Spring mean Temperature (˚C)  Summer mean Temperature (˚C)  Autumn mean Temperature (˚C)  Winter Precipitation (mm)  Spring Precipitation (mm)  Summer Precipitation (mm)  Autumn Precipitation (mm) | January – December  mean Temperatures (˚C)  January – December maximum Temperatures (˚C)  January – December minimum Temperatures (˚C)  January – December Precipitation (mm) |

### **Appendix S4** – Proportion of environmental differences among populations explained by the first PC axis (PC1), the second PC axis (PC2), and the combination of both PC axes (PC1 and PC2) for 15 plant species.

| Species | Variance explained by PC1 (%) | Variance explained by PC2 (%) | Variance explained by PC1 and PC2 (%) |
| --- | --- | --- | --- |
| *Betula maximowicziana* | 62.5 | 20.7 | 83.2 |
| *Centaurea solstitialis* | 52.9 | 21.8 | 74.7 |
| *Helianthus annuus* | 77.7 | 12.8 | 90.5 |
| *Helianthus argophyllus* | 53.6 | 38.6 | 92.2 |
| *Mimulus guttatus* | 65.7 | 21.7 | 87.4 |
| *Mimulus lacinatus* | 80.2 | 11.6 | 91.8 |
| *Narcissus papyraceus* | 53 | 28.7 | 81.7 |
| *Nothofagus alpina* | 65 | 24.1 | 89.1 |
| *Nothofagus glauca* | 54.7 | 27.4 | 82.1 |
| *Nothofagus obliqua* | 59.8 | 28.7 | 88.5 |
| *Picea sitchensis* | 75 | 18 | 93 |
| *Populus balsamifera* | 61.8 | 20.2 | 82 |
| *Populus tremula* | 75.7 | 16.1 | 91.8 |
| *Rhododendron oldhamii* | 66.1 | 19.4 | 85.5 |
| *Shorea leprosula* | 54.7 | 19.2 | 73.9 |

### Appendix S5 – Scatterplots of the relationship between genetic differentiation, based on Nei’s F_ST_, and environmental distance (red), genetic differentiation and geographic distance (blue) as well as between environmental distance and geographic distance (black) for all 19 genetic and genomic datasets considered in this study. Geographic distances were measured as the Euclidean distance between GPS coordinates of sampled populations (latitude, longitude), whereas environmental distances were measured as the Euclidean distance between the two major environmental principal components associated with populations. At the top of each plot figure the correlation coefficient (r) between distances assessed and its significance statistic (P), both estimated using a mantel test.

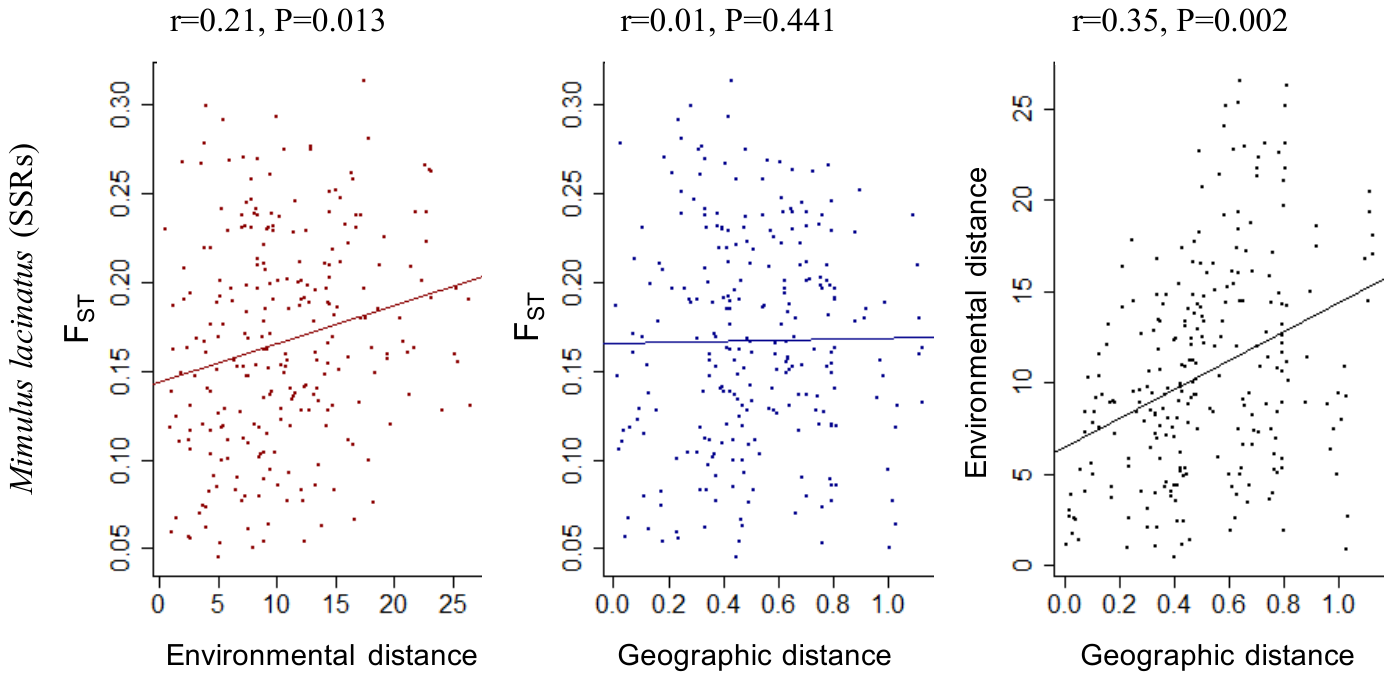

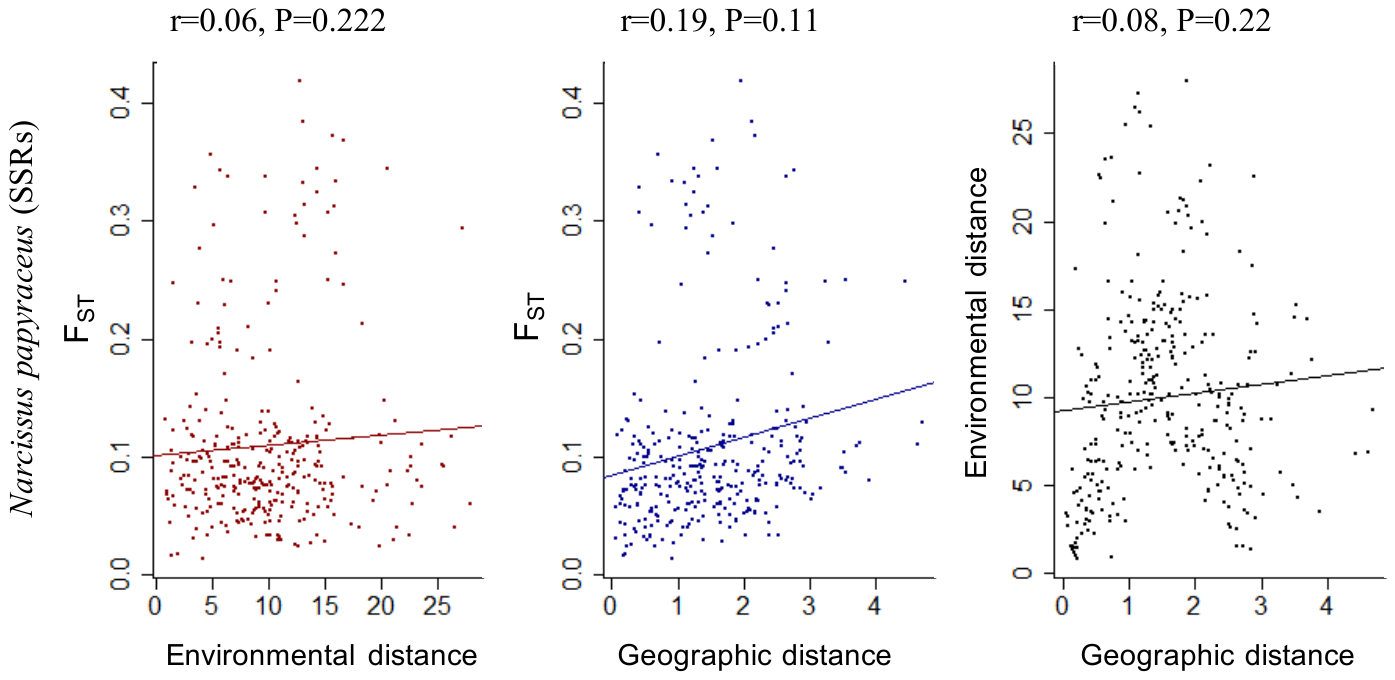

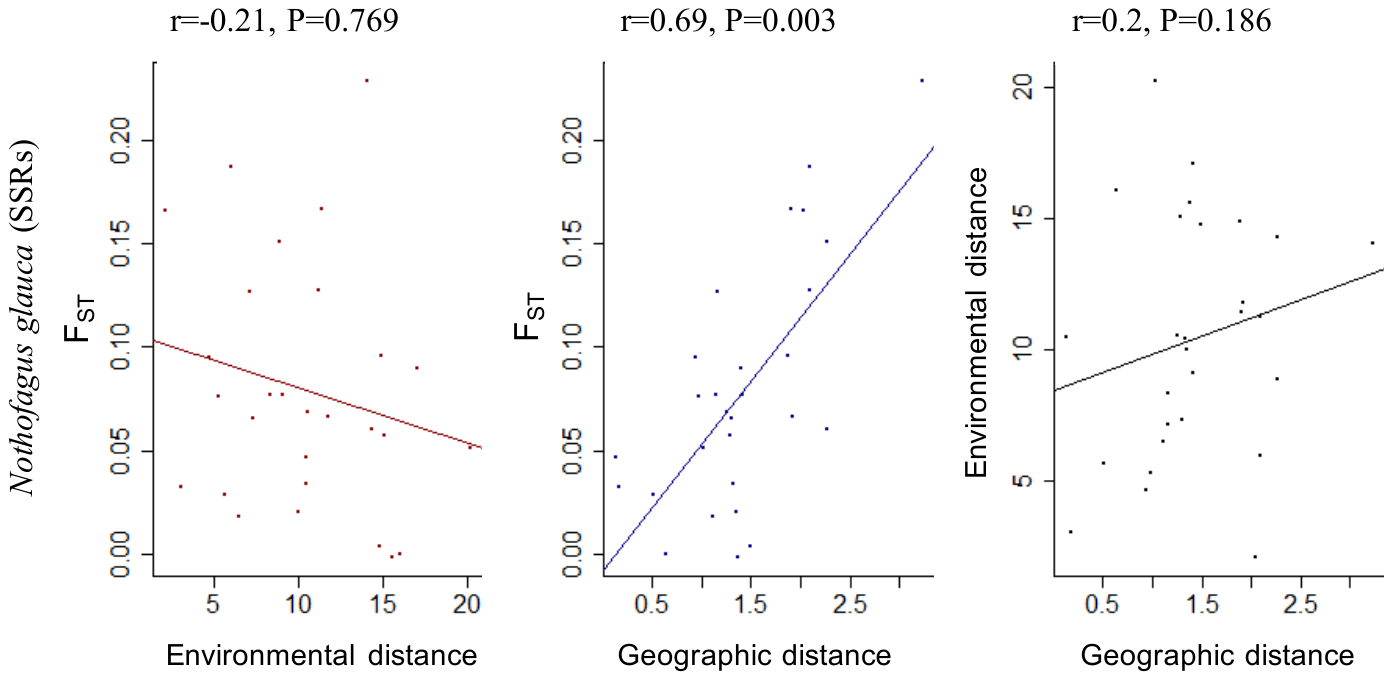

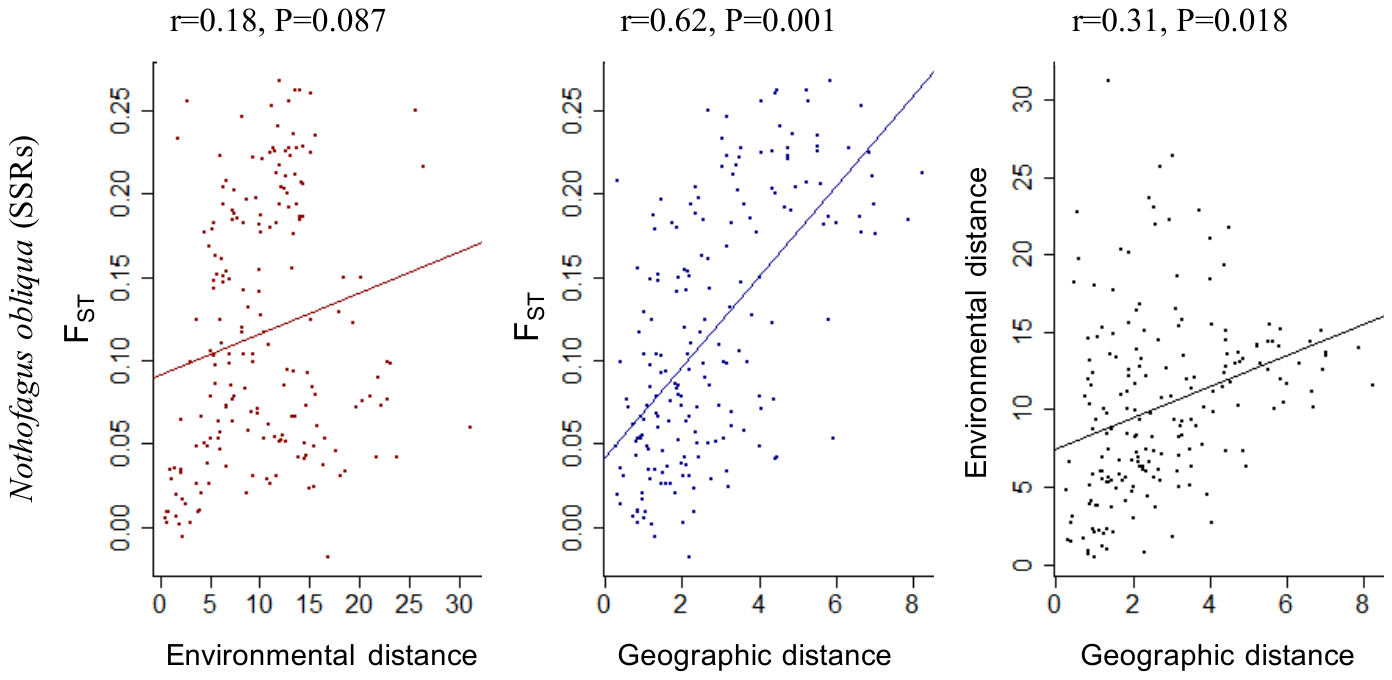

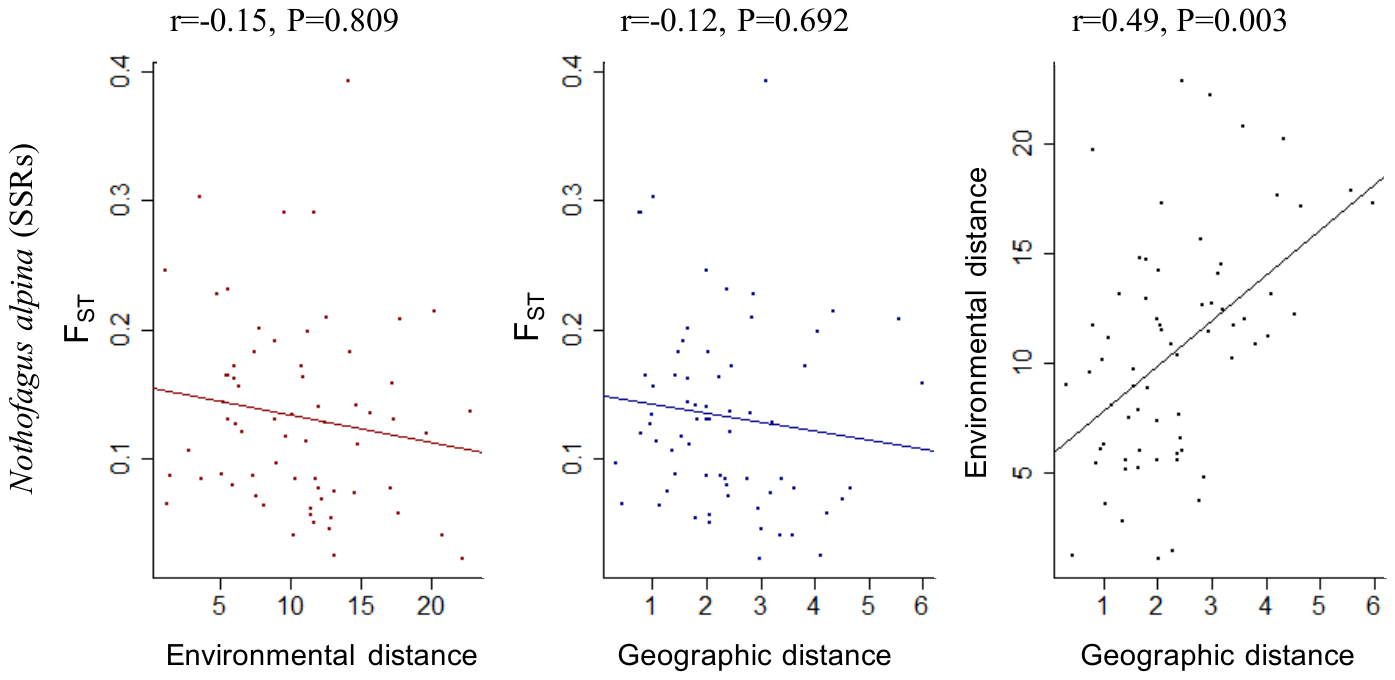

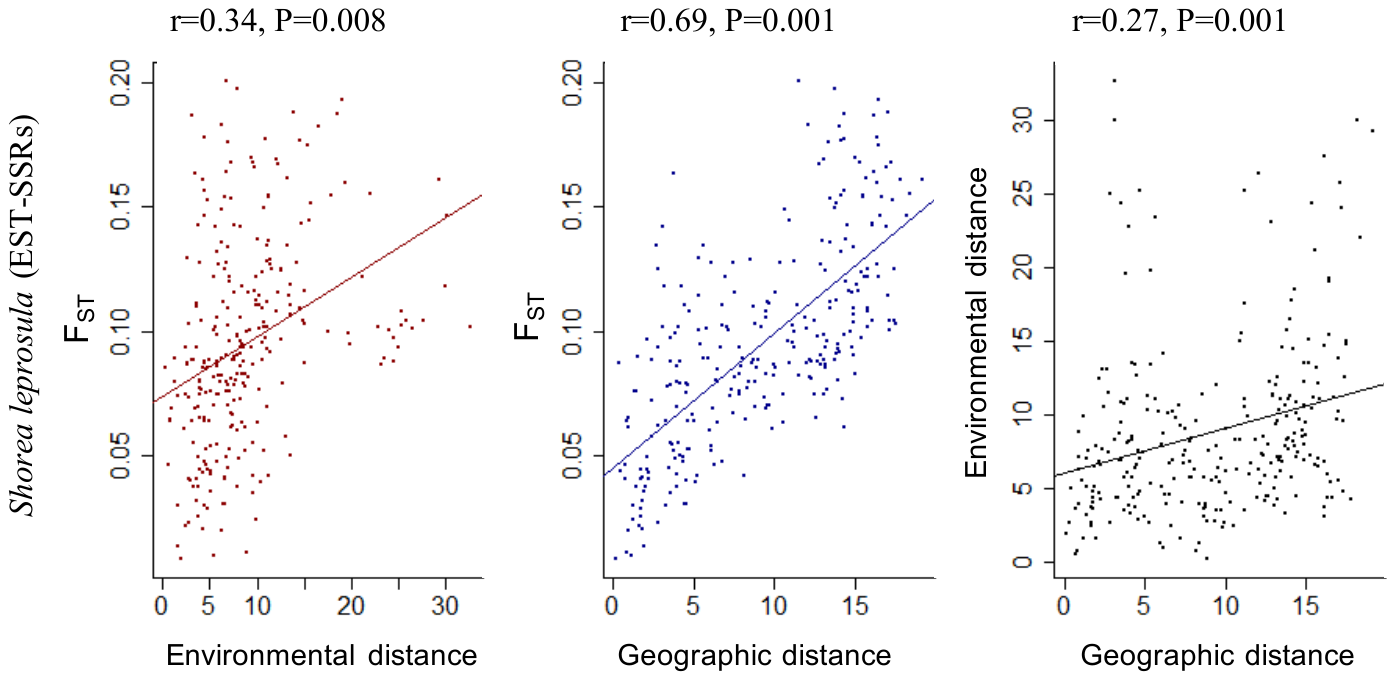

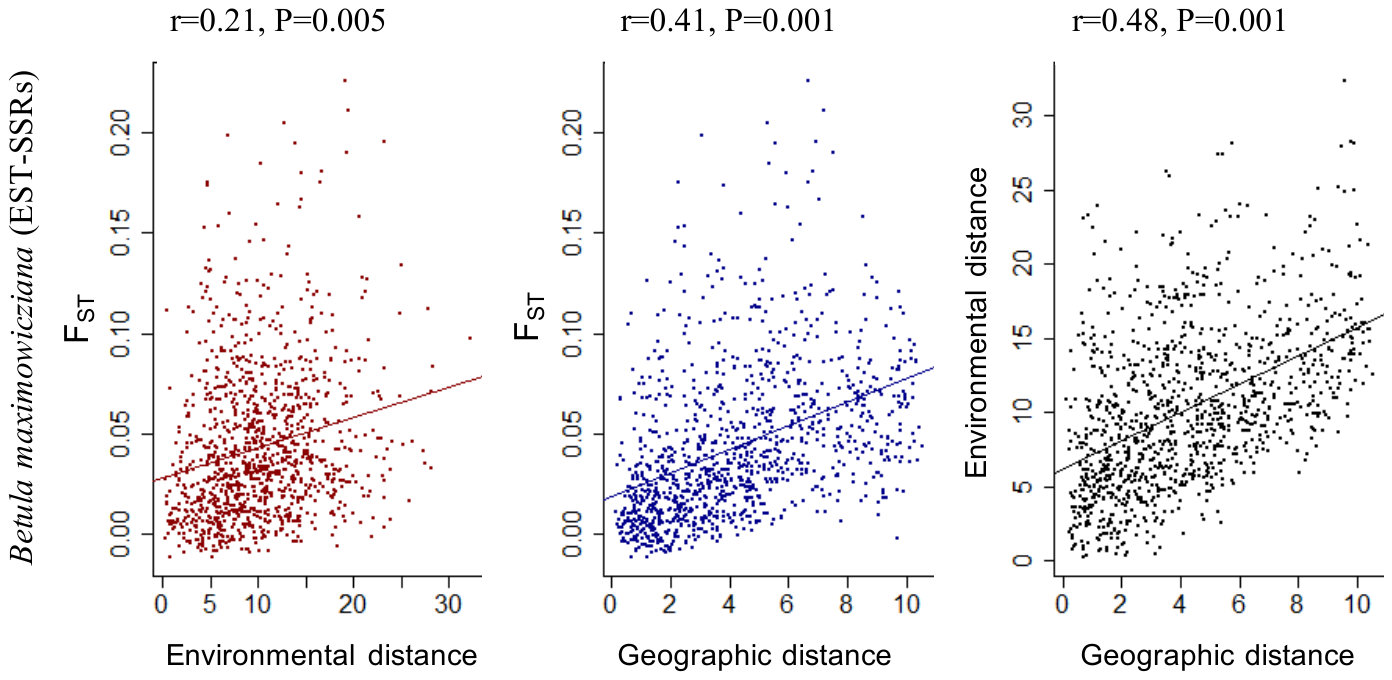

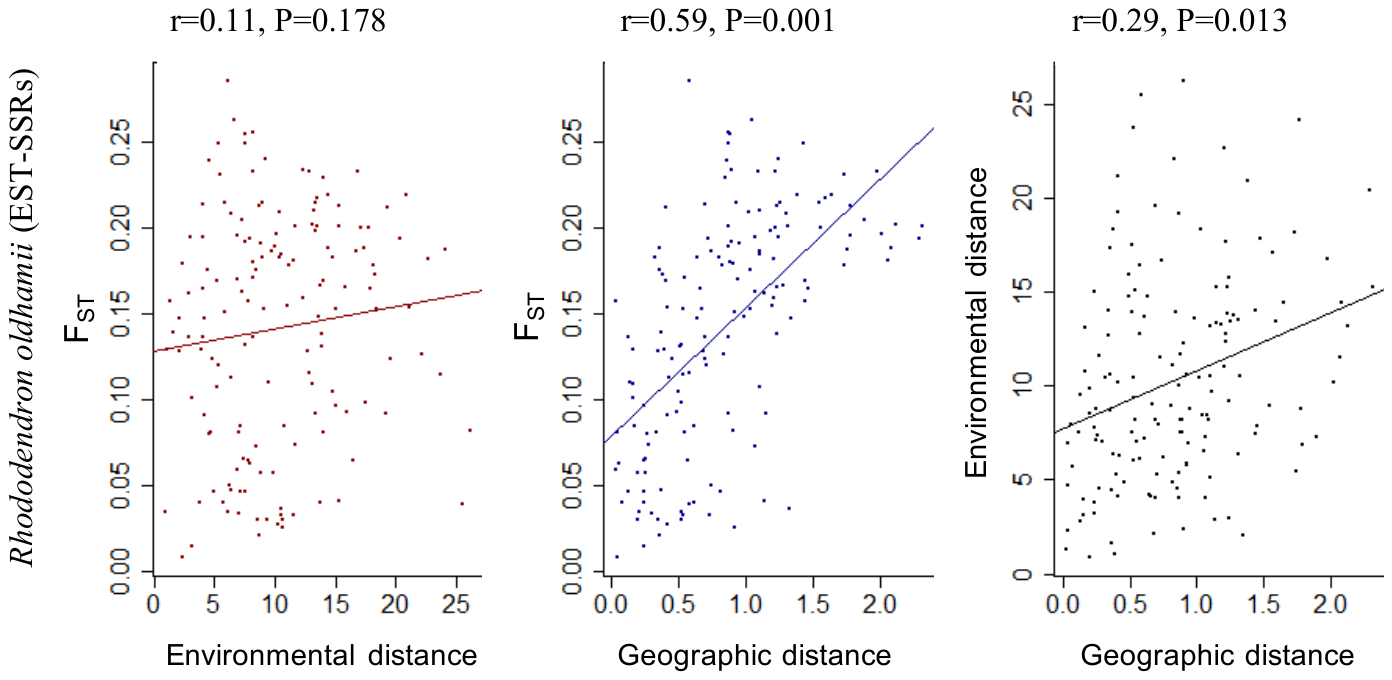

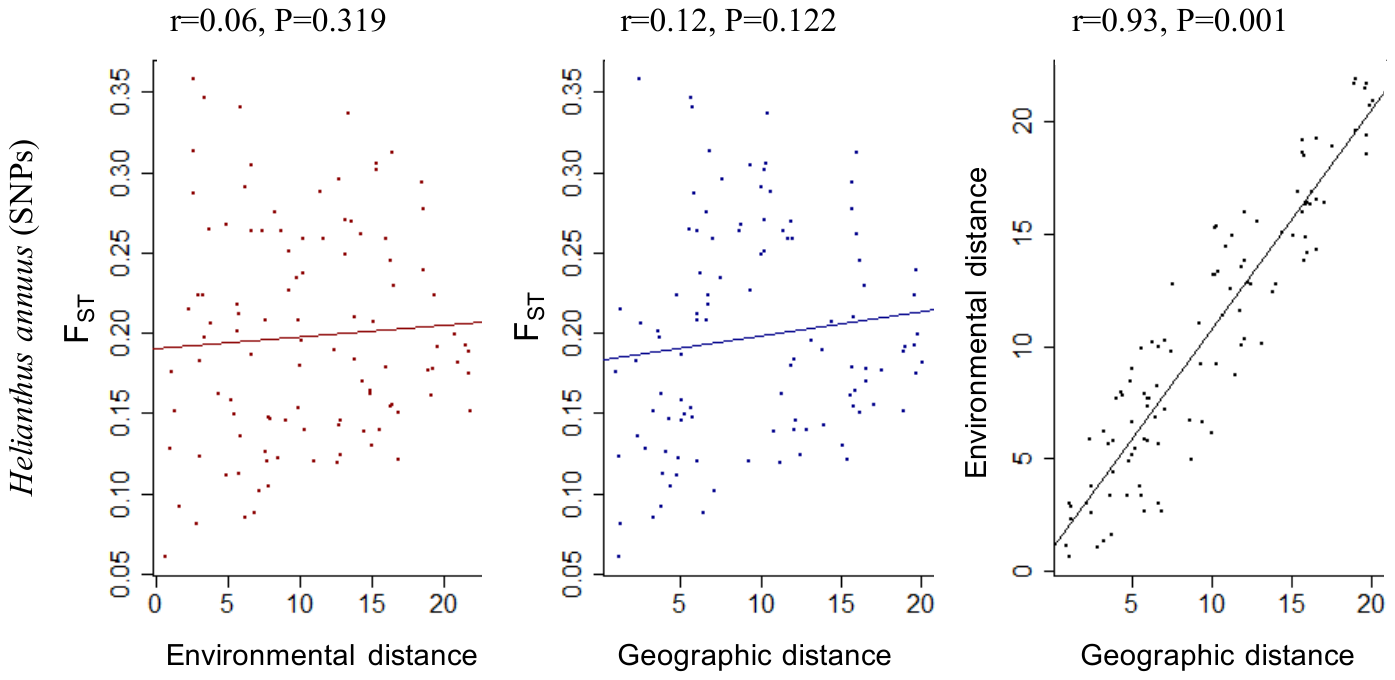

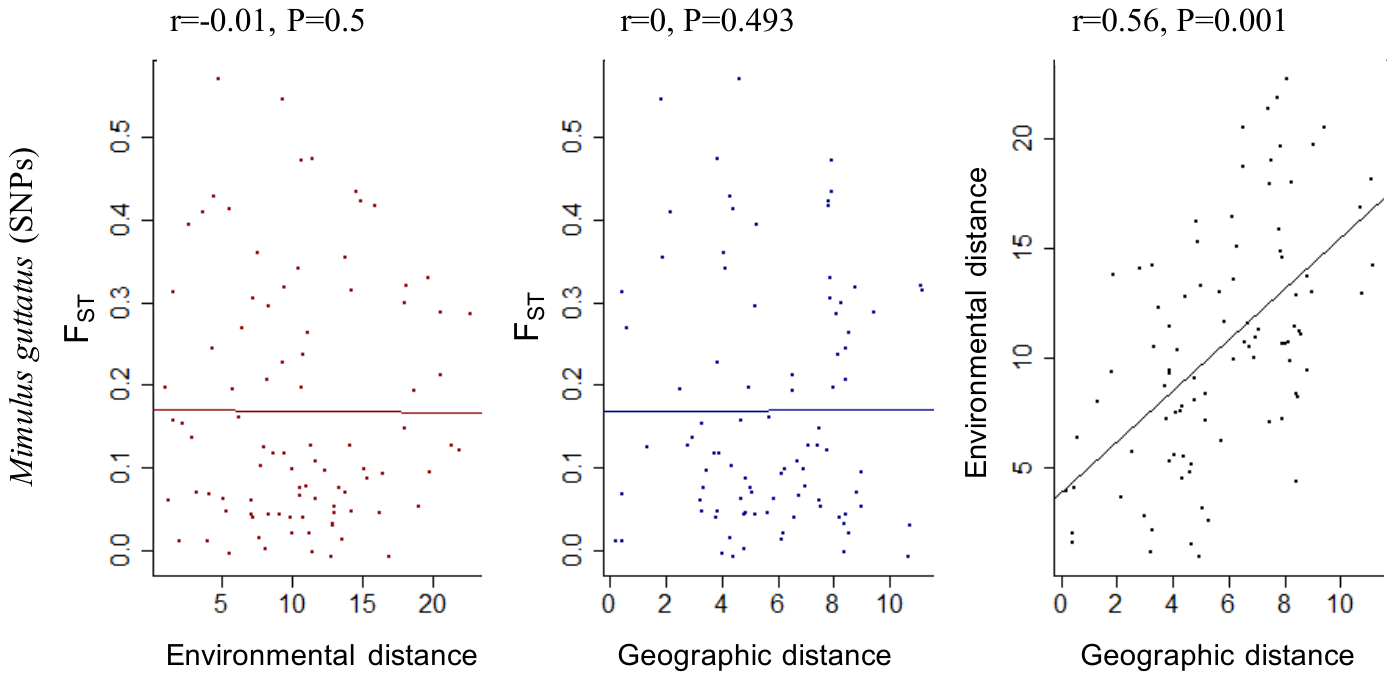

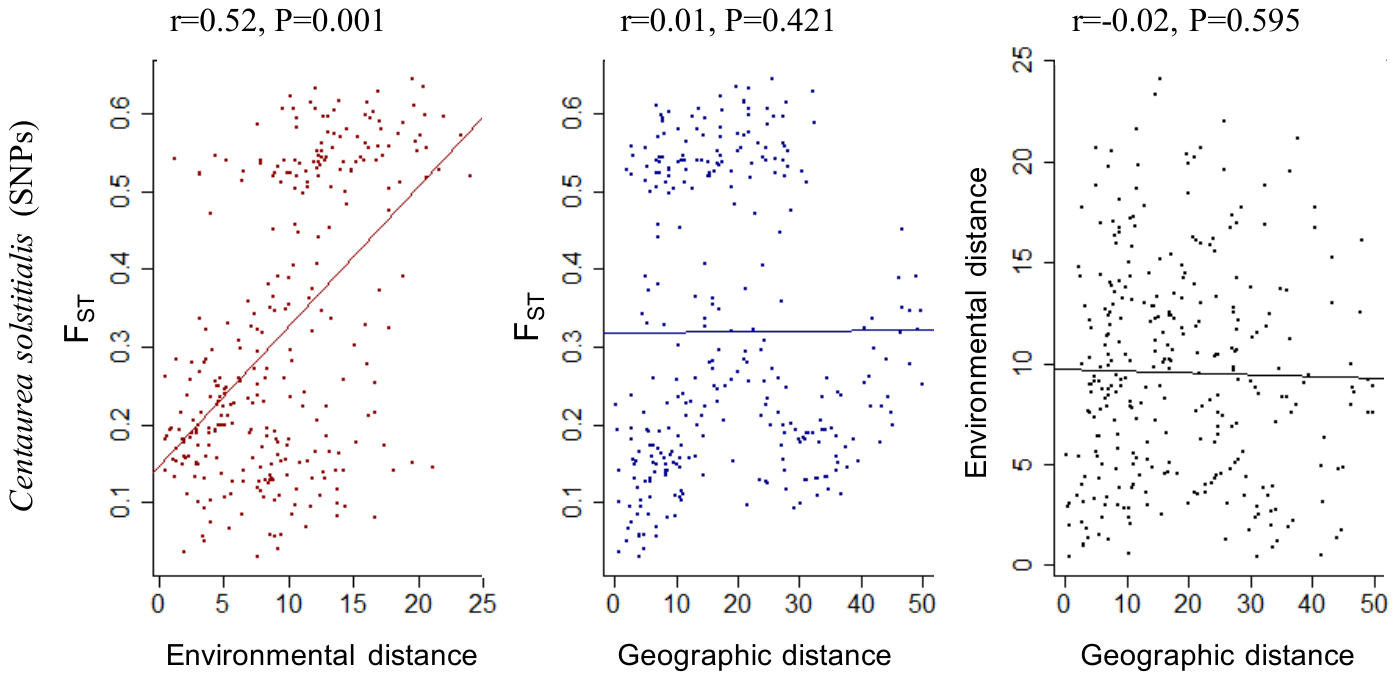

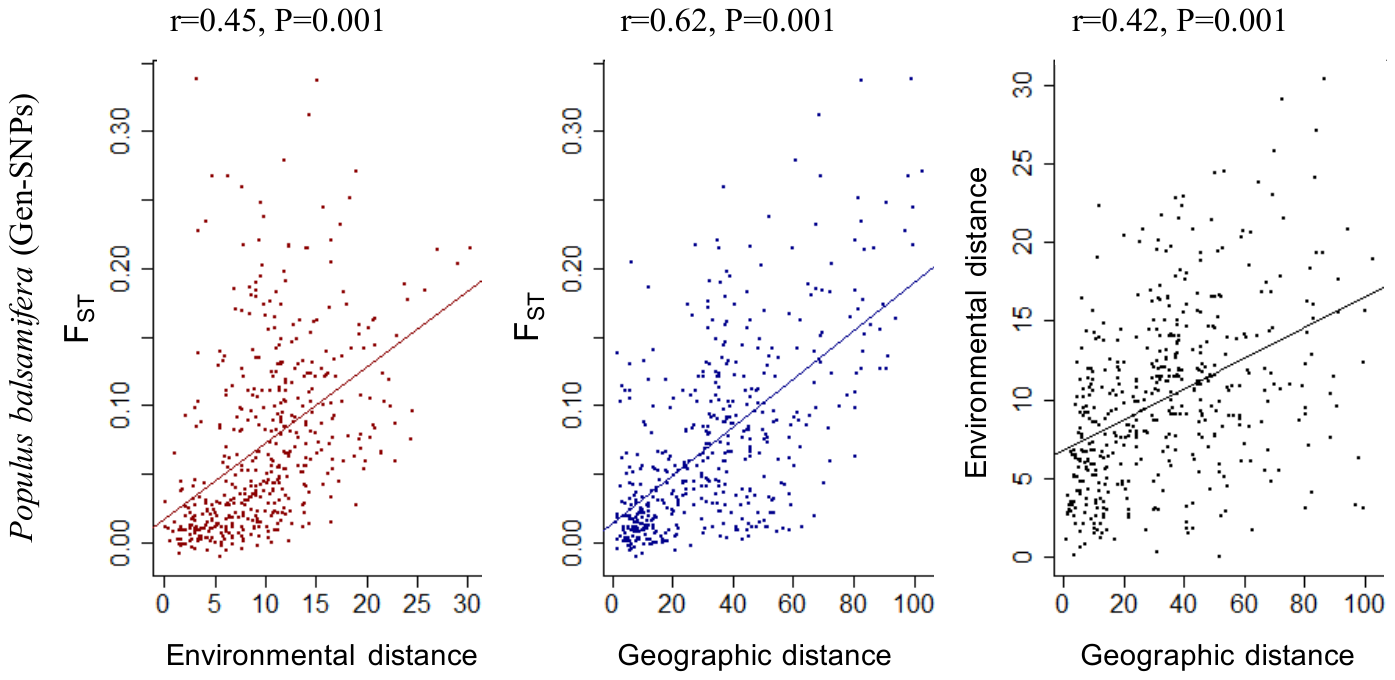

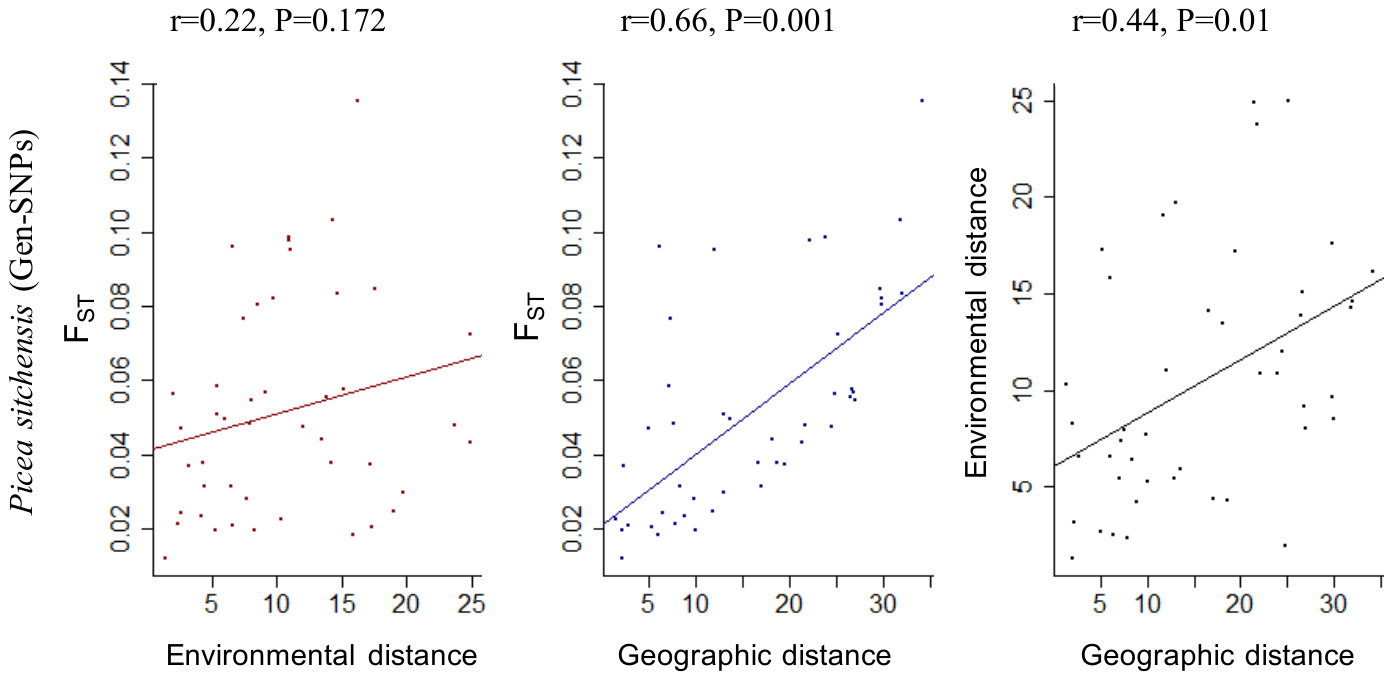

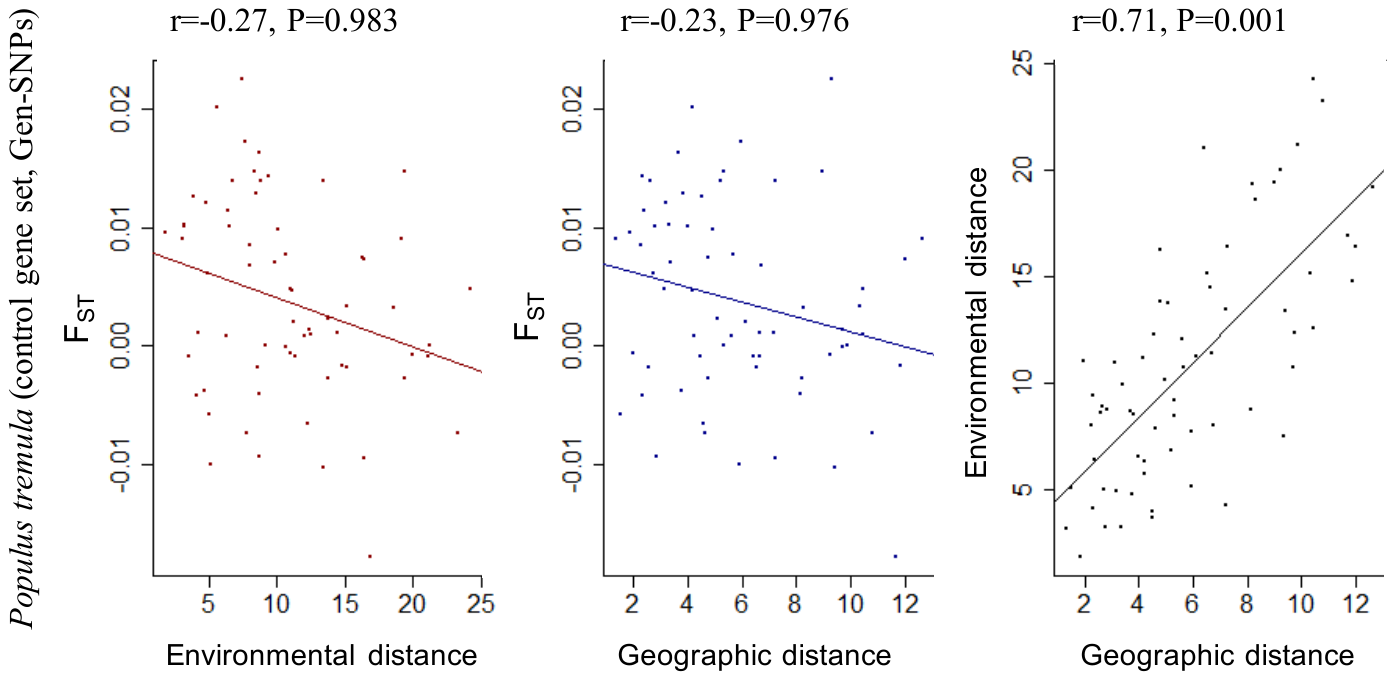

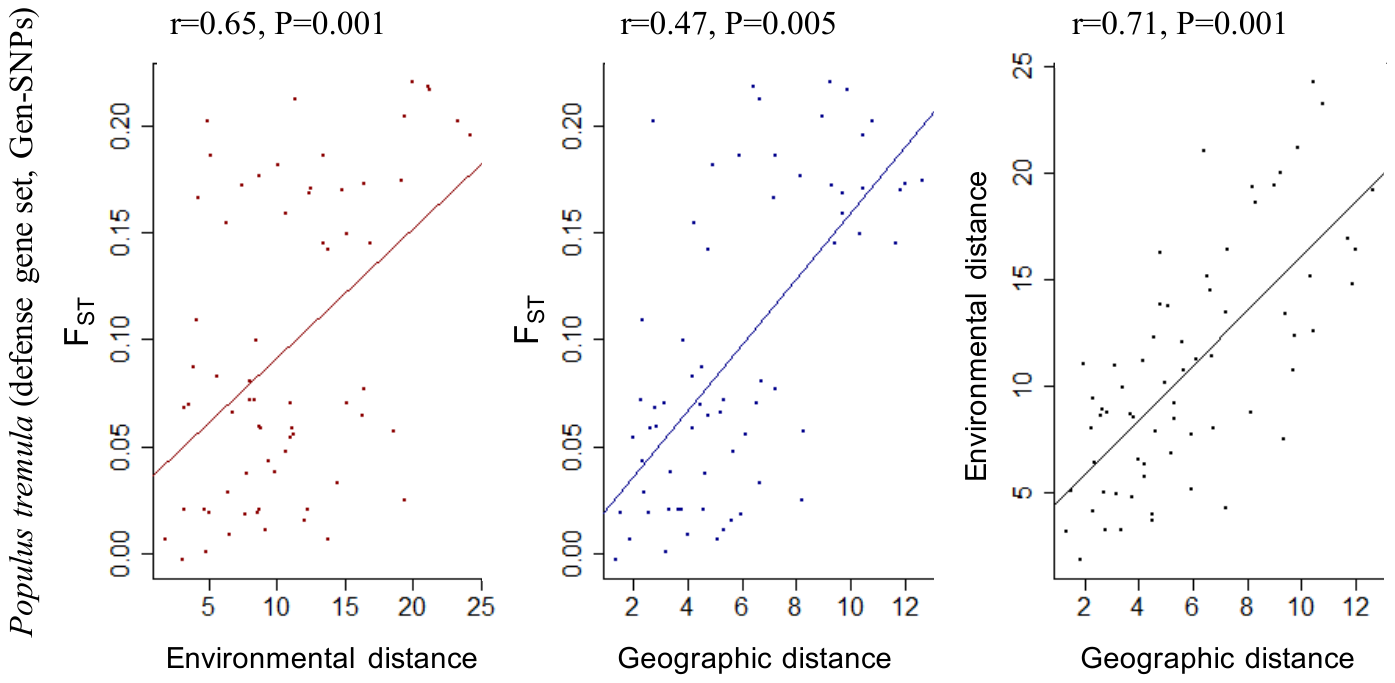

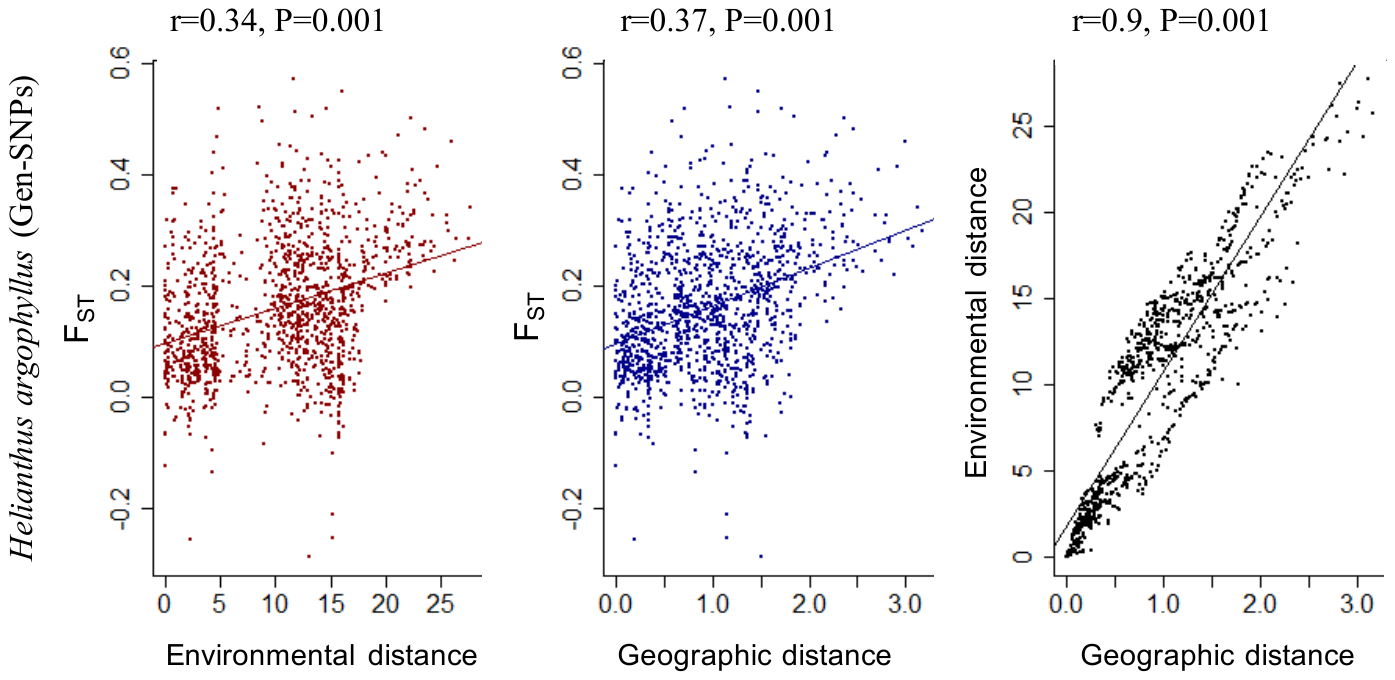

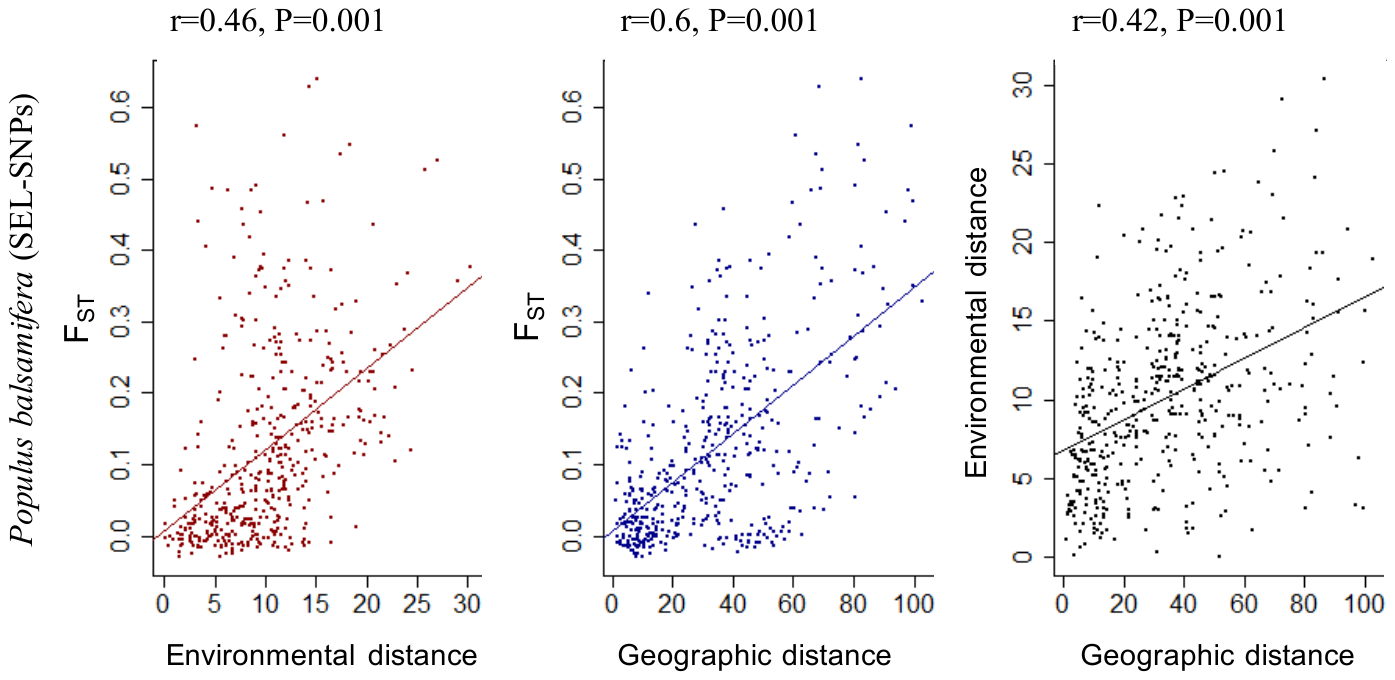

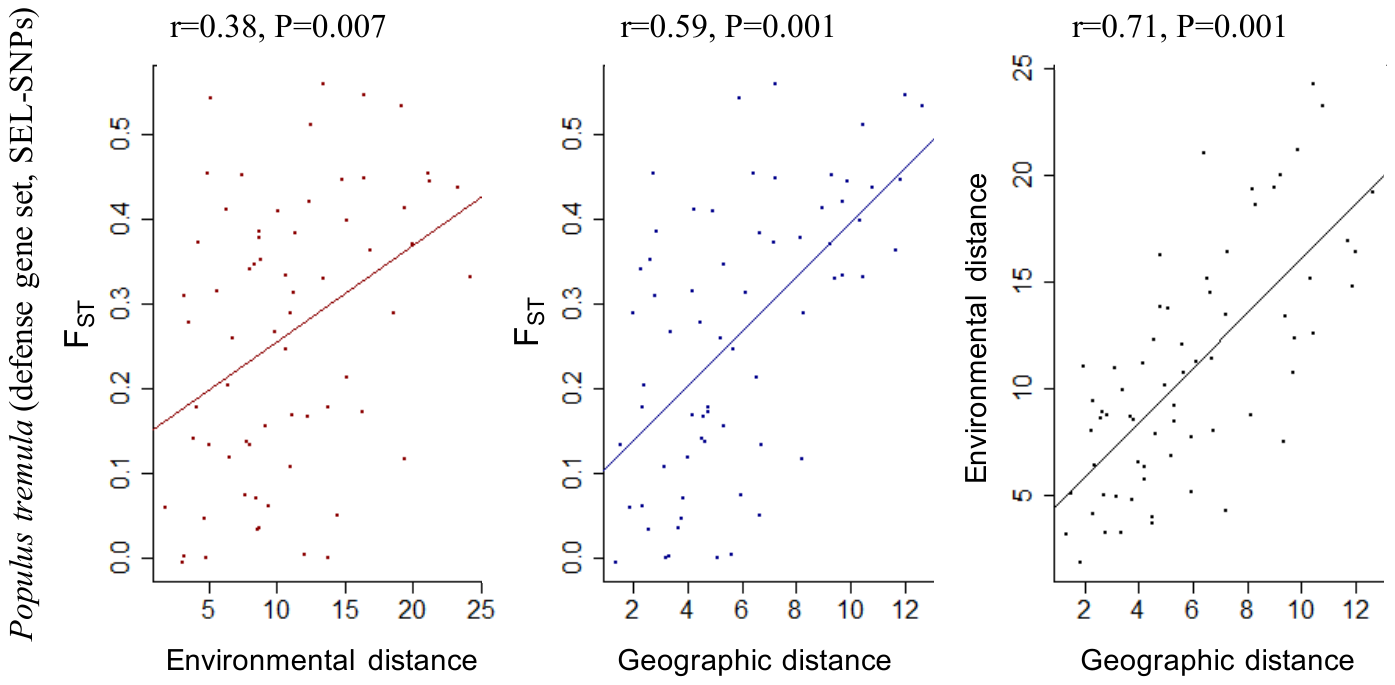

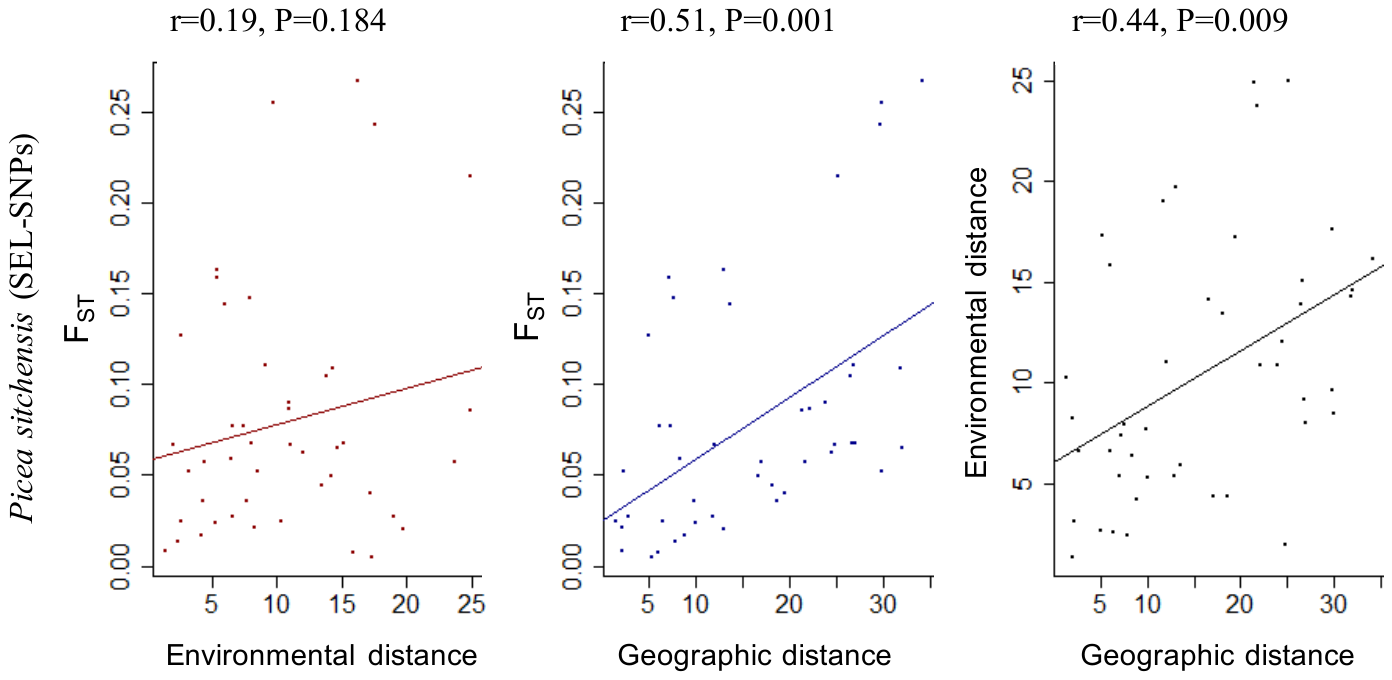

### Appendix S6 – Proportion of allelic diversity captured (y-axis) within a population (x-axis; one point illustrates one population) when *N* individuals were randomly sampled. Different colors represent different datasets. The legend indicates which color is associated with which dataset and provides the value of *N* used for simulations in brackets. Dashed red lines represent the threshold above which 80% or more of allelic diversity is captured and error bars represent Student 95% confidence interval calculated from 500 iterations. (a) Datasets included in “idealized” and “realistic” simulations. For each dataset, *N* = *N_80%_*. (b) Datasets discarded from “idealized” and “realistic” simulations. *N* represents the size of the smallest population within each dataset.

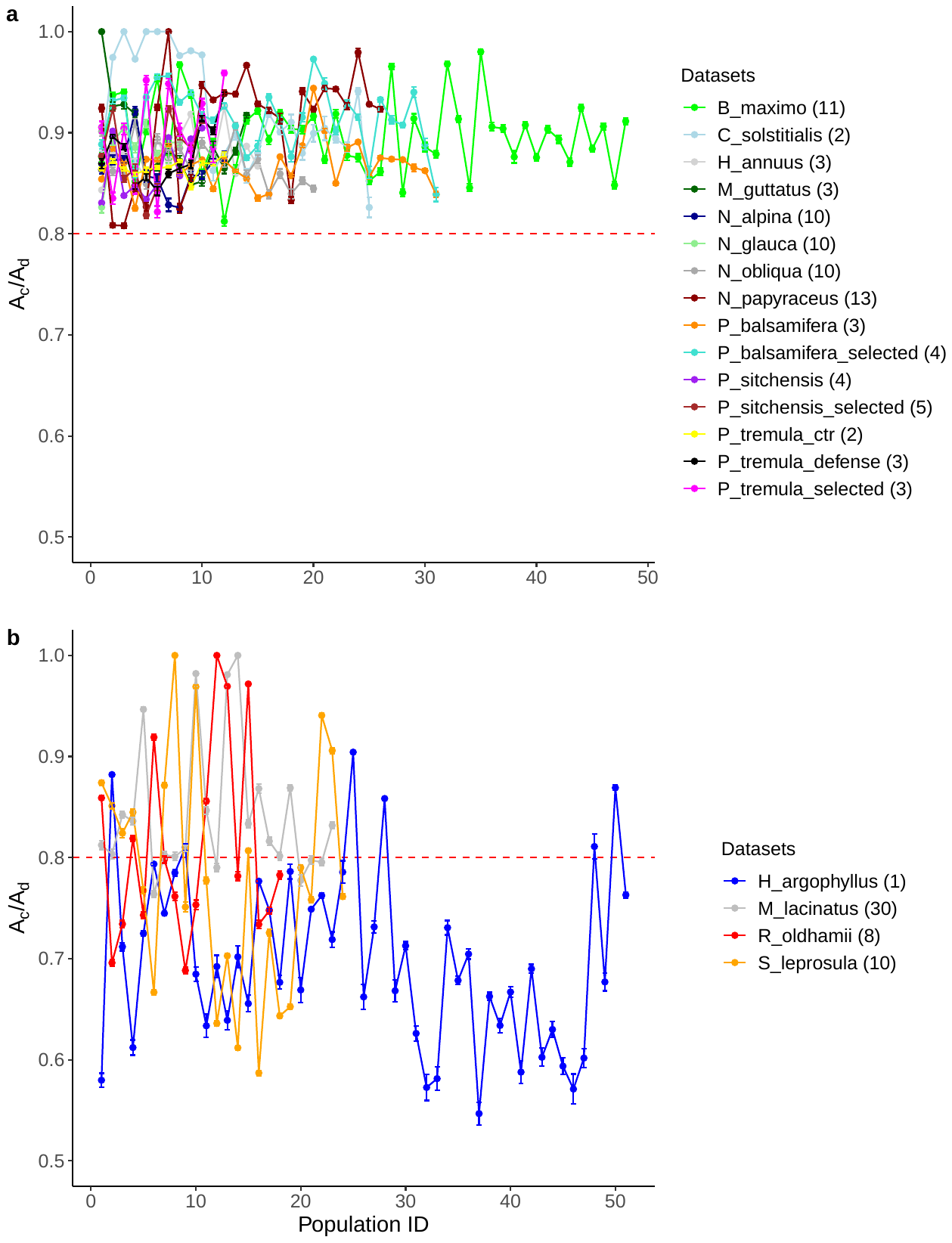

### **Appendix S7** – List of parameters tested and used for realistic and idealized simulations given per dataset.

| Species |  | Genetic marker  (Number of loci) |  | Number of populations |  | Number of individuals per population | | |  | N_80%_ |  | Np | | | |
| --- | --- | --- | --- | --- | --- | --- | --- | --- | --- | --- | --- | --- | --- | --- | --- |
|  |  |  |  |  |  | Min | Max | Mean (SD) |  |  |  | 30-40% | 50-60% | 70-80% | 90-100% |
| *Betula maximowicziana* |  | EST-SSRs (12) |  | 48 |  | 21 | 49 | 29.5 (4.8) |  | 11 |  | 15 | 24 | 36 | 48 |
| *Centaurea solstitialis* |  | SNPs (747) |  | 25 |  | 2 | 19 | 9 (6.2) |  | 2 |  | 8 | 13 | 19 | 25 |
| *Helianthus annuus* |  | SNPs (246) |  | 15 |  | 12 | 20 | 19.1 (2.3) |  | 3 |  | 5 | 8 | 11 | 15 |
| *Helianthus argophyllus* |  | Gen-SNPs (68) |  | 51 |  | 1 | 29 | 10.9 (9.7) |  | NA |  | NA | NA | NA | NA |
| *Mimulus guttatus* |  | SNPs (62) |  | 14 |  | 10 | 21 | 18.6 (2.6) |  | 3 |  | 5 | 7 | 11 | 14 |
| *Mimulus lacinatus* |  | SSRs (8) |  | 23 |  | 30 | 49 | 41.3 (5.5) |  | NA |  | NA | NA | NA | NA |
| *Narcissus papyraceus* |  | SSRs (8) |  | 26 |  | 13 | 20 | 16.2 (2.2) |  | 13 |  | 8 | 13 | 20 | 26 |
| *Nothofagus alpina* |  | SSRs (7) |  | 12 |  | 16 | 16 | 16 (0) |  | 10 |  | 4 | 6 | 9 | 12 |
| *Nothofagus glauca* |  | SSRs (7) |  | 8 |  | 16 | 16 | 16 (0) |  | 10 |  | 3 | 4 | 6 | 8 |
| *Nothofagus obliqua* |  | SSRs (6) |  | 20 |  | 16 | 16 | 16 (0) |  | 10 |  | 6 | 10 | 15 | 20 |
| *Picea sitchensis* |  | Gen-SNPs (339) |  | 10 |  | 12 | 46 | 28.6 (13.2) |  | 4 |  | 3 | 5 | 8 | 10 |
|  |  | SEL-SNPs (34) |  | 10 |  | 12 | 46 | 28.6 (13.2) |  | 5 |  | 3 | 5 | 8 | 10 |
| *Populus balsamifera* |  | Gen-SNPs (284) |  | 31 |  | 10 | 18 | 14.3 (1.4) |  | 3 |  | 10 | 16 | 23 | 31 |
|  |  | SEL-SNPs (33) |  | 31 |  | 10 | 18 | 14.3 (1.4) |  | 4 |  | 10 | 16 | 23 | 31 |
| *Populus tremula* |  | Gen-SNPs (93)  [control set] |  | 12 |  | 6 | 10 | 9.6 (1.2) |  | 2 |  | 4 | 6 | 9 | 12 |
|  |  | Gen-SNPs (71)  [defense set] |  | 12 |  | 6 | 10 | 9.2 (1.1) |  | 3 |  | 4 | 6 | 9 | 12 |
|  |  | SEL-SNPs (10) |  | 12 |  | 6 | 10 | 9.2 (1.1) |  | 3 |  | 4 | 6 | 9 | 12 |
| *Rhododendron oldhamii* |  | EST-SSRs (26) |  | 18 |  | 8 | 31 | 18.7 (6.9) |  | NA |  | NA | NA | NA | NA |
| *Shorea leprosula* |  | EST-SSRs (27) |  | 24 |  | 10 | 59 | 30.25 (16.1) |  | NA |  | NA | NA | NA | NA |

### Appendix S8 – Regression statistics of genetic parameters assessed (F_ST_ and A_c_/A_d_ ; see Fig. 2 for details) separated first by comparisons (Env-Rand, Geo-Rand, Env & Geo-Rand) and then by within-population sampling scenarios (realistic vs idealized). For each genetic parameter, slopes and SEs are given per genetic marker class.

| Genetic marker class | Env-Rand | |  | Geo-Rand | |  | Env & Geo-Rand | |
| --- | --- | --- | --- | --- | --- | --- | --- | --- |
|  | realistic  slope (SE) | idealized  slope (SE) |  | realistic  slope (SE) | idealized  slope (SE) |  | realistic  slope (SE) | idealized  slope (SE) |
| *F_ST_* |  |  |  |  |  |  |  |  |
| Neutral | -0.005$($0.004) | -0.006$($0.004) |  | 0.003$($0.007) | 0.002$($0.007) |  | -0.001$($0.005) | -0.001$($0.005) |
| Functional | -0.004$($0.002)^*^ | -0.004$($0.002)^*^ |  | -0.008$($0.004) | -0.008$($0.004) |  | -0.006$($0.003)^*^ | -0.006$($0.003)^*^ |
| Adaptive | -0.024$($0.006)^*^ | -0.023$($0.006)^*^ |  | -0.035$($0.006)^*^ | -0.035$($0.006)^*^ |  | -0.023$($0.001)^*^ | -0.022$($0.002)^*^ |
| *A_c_/A_d_* |  |  |  |  |  |  |  |  |
| Neutral | -0.001(0.007) | 0.001(0.007) |  | 0$($0.006) | 0.001$($0.007) |  | -0.005$($0.005) | -0.005$($0.006) |
| Functional | -0.003(0.001)^*^ | -0.006$($0.004) |  | 0.001$($0.003) | 0.005$($0.004) |  | -0.001$($0.002) | -0.003$($0.004) |
| Adaptive | -0.008$($0.004) | -0.006$($0.002)^*^ |  | -0.005$($0.005) | 0$($0.005) |  | -0.01$($0.005) | -0.006$($0.002)^*^ |

* indicate that observed slopes are significantly different from zero (α=0.05).
